# Characterization of tumor heterogeneity through segmentation-free representation learning

**DOI:** 10.1101/2024.09.05.611431

**Authors:** Jimin Tan, Hortense Le, Jiehui Deng, Yingzhuo Liu, Yuan Hao, Michelle Hollenberg, Wenke Liu, Joshua M. Wang, Bo Xia, Sitharam Ramaswami, Valeria Mezzano, Cynthia Loomis, Nina Murrell, Andre L. Moreira, Kyunghyun Cho, Harvey Pass, Kwok-Kin Wong, Yi Ban, Benjamin G. Neel, Aristotelis Tsirigos, David Fenyö

## Abstract

The interaction between tumors and their microenvironment is complex and heterogeneous. Recent developments in high-dimensional multiplexed imaging have revealed the spatial organization of tumor tissues at the molecular level. However, the discovery and thorough characterization of the tumor microenvironment (TME) remains challenging due to the scale and complexity of the images. Here, we propose a self-supervised representation learning framework, CANVAS, that enables discovery of novel types of TMEs. CANVAS is a vision transformer that directly takes high-dimensional multiplexed images and is trained using self-supervised masked image modeling. In contrast to traditional spatial analysis approaches which rely on cell segmentations, CANVAS is segmentation-free, utilizes pixel-level information, and retains local morphology and biomarker distribution information. This approach allows the model to distinguish subtle morphological differences, leading to precise separation and characterization of distinct TME signatures. We applied CANVAS to a lung tumor dataset and identified and validated a monocytic signature that is associated with poor prognosis.

## Introduction

Tumor heterogeneity, characterized by variations in cellular and molecular features such as genetic, epigenetic, and phenotypic profiles, presents a significant challenge to effective treatment. In addition to cancer cell intrinsic hallmarks, including sustained proliferation, evading growth suppressors, resisting cell death, invasion and metastasis, recent emerging hallmarks, including tumor-promoting inflammation and evading immune destruction that come from a repertoire of recruited normal cells generates additional complexity and therapeutic vulnerability for cancer^1,2^ . This variability within a single tumor is characterized by the tumor microenvironment (TME), which consists of immune cells, fibroblasts, neovasculature, and extracellular matrix. The TME can influence tumor cells by facilitating nutrient supply and immune evasion, thereby contributing to increased heterogeneity and uncontrolled growth of malignant cancer cells that are resistant to cancer-cell intrinsic targeted therapy and chemotherapy. The distinct metabolic gradients within the TME can create ecological niches that sustain different tumor subpopulations^3,4^ . Understanding the role of TME not only led to breakthrough therapies such as immunotherapy drugs, but also suggests future strategies towards curing cancer^5–9^.

Recent advances in spatial omics have enabled the thorough investigation of TMEs^10–18^. However, the vast amount of data generated by high-dimensional spatial imaging require new high-throughput analysis methods. A typical preprocessing method to reduce spatial dimensionality of the data is to perform cell segmentations and to consider the cells as the basic units for the analysis. By treating cells as nodes and their adjacency as edges, spatial imaging data can be reformulated as graphs. Subsequently, various graph-based operations and modeling can be performed on the imaging data^19–24^.

However, c ell segmentation-based approaches simplify analysis but are prone to bias from external cell segmentation tools. More importantly, cell segmentation discards crucial morphological features that potentially represent the underlying pathology^25,26^. To capture the details of the imaging data without bias, we argue that a method needs to efficiently encode pixel level information. This task is challenging for three reasons. First, a dedicated model is required to learn the features from scratch, as no external annotation can be leveraged. Second, the model must have high capacity to effectively recognize the heterogeneity of TMEs. Third, a high-throughput method is required due to the large size of spatial omics data.

Here, we propose a novel analysis framework CANVAS (CANcer Vision AutoencoderS), a high-capacity deep neural network that addresses these challenges. CANVAS is trained by self-supervised learning without requiring labels, annotations, or segmentation. The model also learns directly from pixel-level multiplexed images and can be trained efficiently on large-scale high-dimensional imaging data.

CANVAS transforms multiplexed tumor images into meaningful representations that allow clustering and characterization of the TMEs. This “end-to-end” design allows CANVAS to learn tissue structure at pixel-level, enables unsupervised discovery of novel TME signatures, and generates new biological and clinical hypotheses. Using CANVAS-derived TME signatures, we show that patients can be categorized into subgroups with distinct CANVAS signatures and clinical outcomes. In addition, we used CANVAS to discover a unique monocytic signature that correlates with poor prognosis, which we validated both in an independent dataset and *in vitro* experiments.

## Results

### Learning a rich representation of the tumor microenvironment

Due to the complexity and diversity of tumor microenvironments, analysis tools that capture protein distributions and their relationship to tissue morphology are crucial for clinical interpretation and biological discovery. For this purpose, w e propose CANVAS, a self-supervised learning framework that performs representation learning on the local tumor microenvironment (Fig. 1). CANVAS is an end-to-end model that learns directly from multiplexed images without requiring cell segmentation, reducing the potential biases stemming from segmentation models and image quality.

**Fig. 1:**
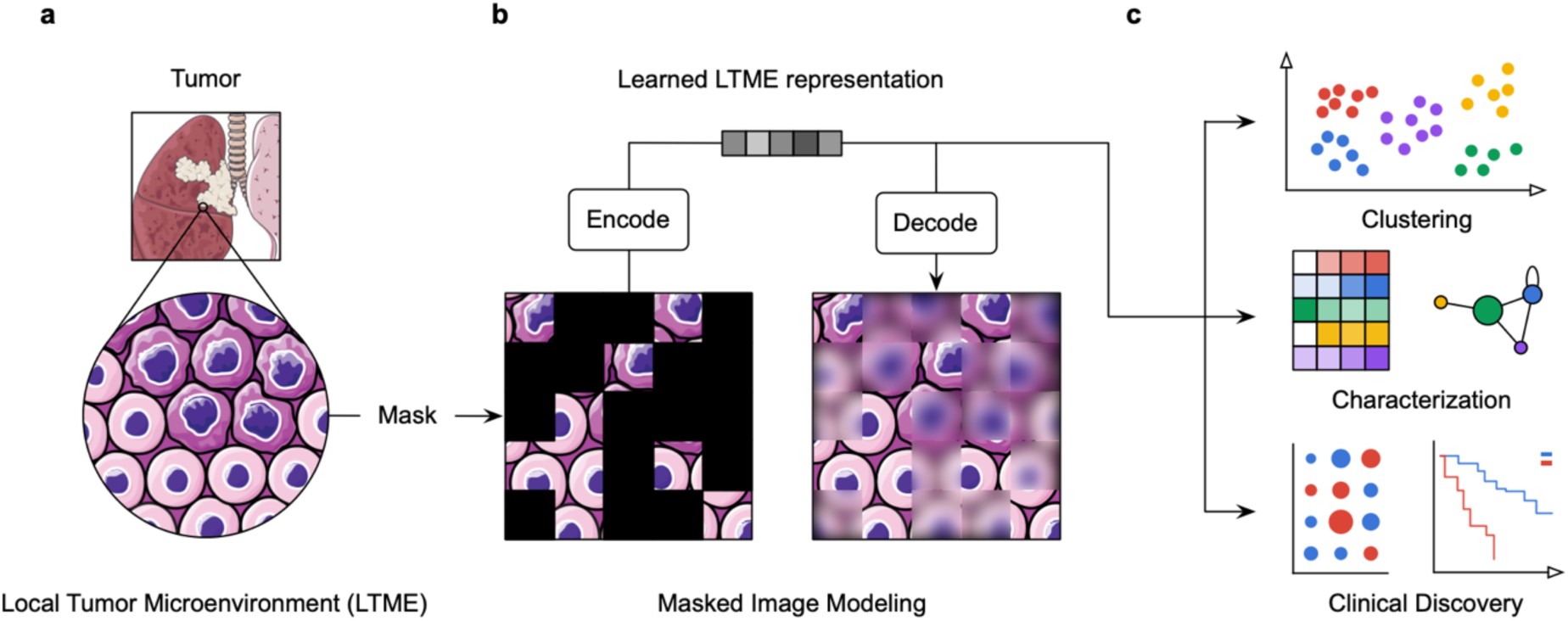
**A schematic of the CANVAS analysis framework. a**, Local tumor microenvironments (LTMEs) are micron scale tissue regions. **b**, CANVAS performs masked image modeling to learn meaningful LTME representations. **c**, Downstream analysis is performed based on the LTME representations generated with the trained encoder without masking.

We applied CANVAS to a lung cancer cohort of 416 patients whose tumors were analyzed using Imaging Mass Cytometry (IMC)^27^ . To study the TME, patient cores were split into small tiles each with a width of 64 micrometers, which we term local tumor microenvironments (LTMEs) (Fig. 2a). The tiling size was chosen to preserve cellular morphology while including around 20 cells to represent cell neighborhoods (see Methods). The LTME images were then normalized and treated as individual training samples for CANVAS.

**Fig. 2:**
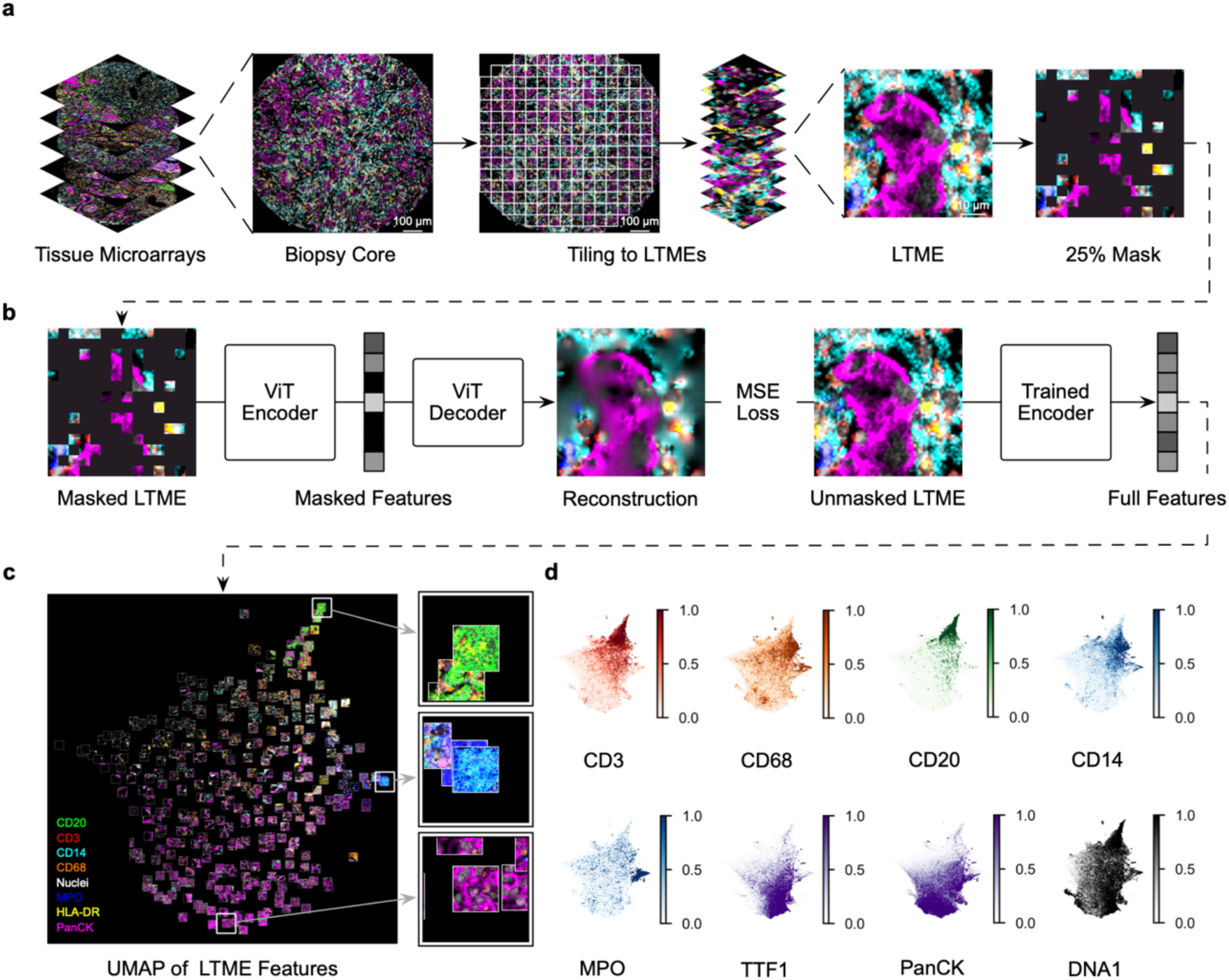
**Analyzing a multiplexed lung IMC dataset with CANVAS. a**, Image preprocessing. Tissue microarray images are separated into LTME tiles. Seventy-five percent (75%) of the image patches and all 18 channels are randomly masked during training. **b**, CANVAS performs masked image modeling on LTMEs. After training the model, the CANVAS encoder uses the full image to generate features for downstream analysis. **c**, Dimensionality reduction with UMAP and visualization of CANVAS features. Each image on the UMAP represents an LTME. LTMEs are colored according to the eight selected markers (bottom left). The right panel section highlights three distinct regions on the UMAP that are enriched in CD20, MPO, and PanCK from top to bottom. **d**, UMAPs colored by different markers.

The CANVAS model consists of an encoder and a decoder, both of which are Vision Transformers (ViTs). ViTs are deep neural network architectures specifically designed for computer vision tasks^28–30^ . In ViTs, images are processed as small patches, each of which is processed independently. This feature gives ViT a unique edge in performing input masking compared with convolutional neural networks (CNNs). This advantage is important, as input masking would leave damaging artifacts in convolution operations.

The input sample is an LTME image with multiple marker channels. The encoder ViT splits this image into multiple 5 by 5-micron patches, and each is considered a token by the transformer model. During training, the encoder processes only 25% of the patches (Fig. 2a). After encoding, dummy tokens representing the missing patches are added to form the joint masked features (Fig. 2b). The decoder is then asked to reconstruct the missing 75% of the patches from the masked features (Fig. 2b). We used channel-wise data augmentation to reduce channel-related bias during training (see Methods).

After training the model, unmasked images were fed into the encoder to produce feature embedding for the LTME (Fig. 2b). We performed dimensionality reduction on the features for all LTMEs and superimposed the LTME images on their UMAP coordinate (see Methods). LTMEs expressing similar markers are enriched in adjacent regions on the UMAP (Fig. 2c, Supplementary Fig. 1). Specifically, the distributions of CD3 (T cells), CD68 (macrophages), CD20 (B cells), CD14 (monocytes), MPO (neutrophils), TTF1 and PanCK (tumor cells), and DNA1 (cell nuclei), are concentrated across the data manifold (Fig. 2d).

### Characterization of LTMEs

Highlighting the individual protein marker is one way to visualize the CANVAS features. Since the model operates at the pixel level, it not only encodes explicit features, including marker intensity, but also implicit features including tissue morphology and cellular interactions. To understand the biological significance of CANVAS features, we performed clustering to group similar features. We defined these clusters as “LTME signatures” and used them for downstream analysis and interpretation (Fig. 3a, see Methods). These signatures can be summarized into broader categories by their marker enrichment (Fig. 3b).

**Fig. 3:**
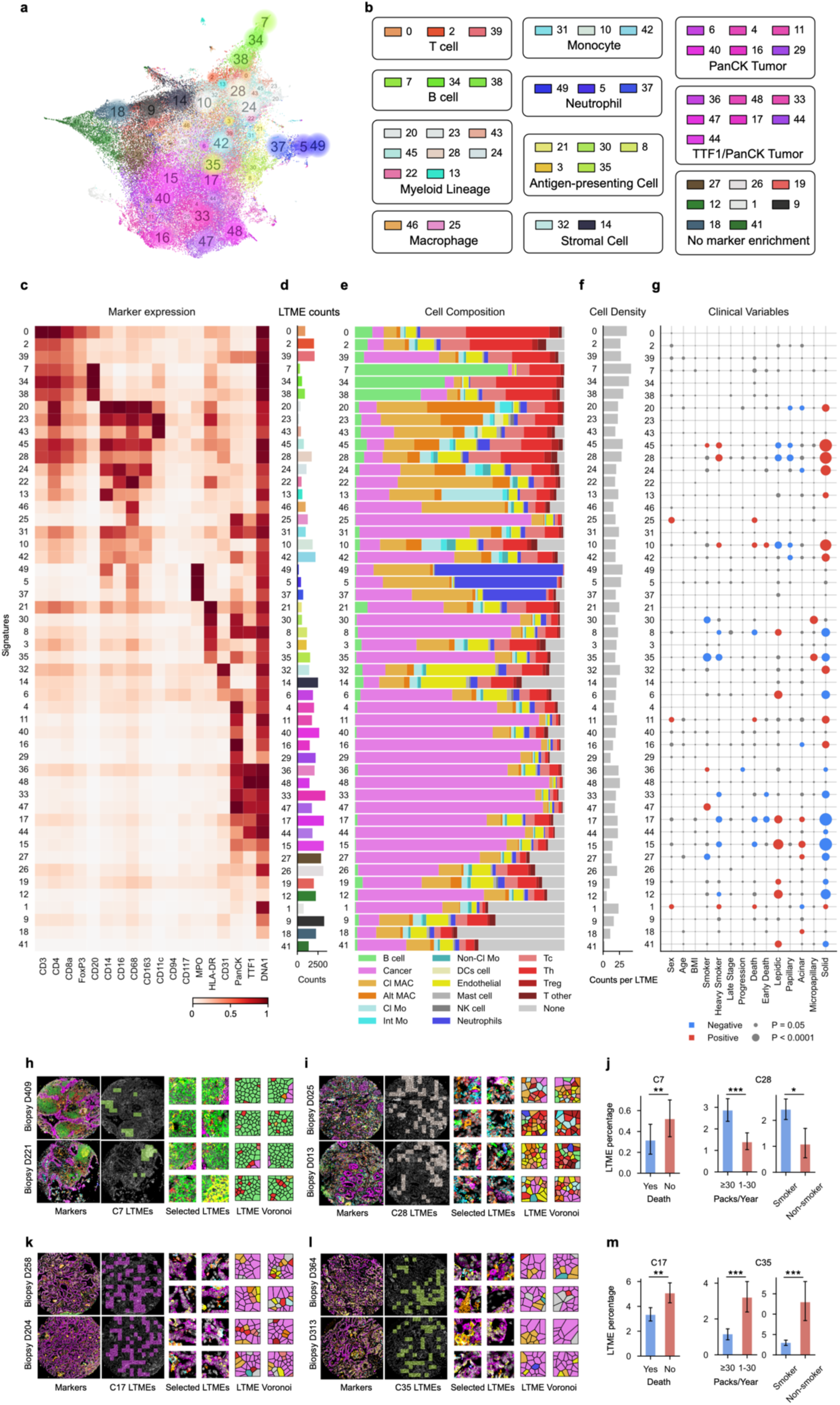
**Characterization of the local tumor microenvironment. a**, Clustering of LTMEs reveals 50 signatures. Signatures are colored according to their marker enrichment and labeled with corresponding dark-colored numbers. Signature colors are consistent across this study. **b**, Grouping of signatures into broader categories. **c**, Marker enrichment by signature. **d**, Distribution of the total number of LTME counts across signatures. **e**, Relative cell type composition by signature. **f**, Average cell counts per LTME for each signature. **g**, Correlation between clinical variables and signatures. Circles represent individual tests correlating clinical variables with signatures. Circle color indicates correlation direction; size denotes statistical significance. Non-significant tests are shown in gray. **h-i**, Selected biopsies enriched in signatures C7 (**h**) and C28 (**i**). The original biopsy is colored with the selected markers from Fig. 2c. Voronoi diagrams are colored by cell based on the code in Fig 3e. **j**, Comparing percentage of signatures C7 and C28 in different clinical conditions. **k-l**, Selected biopsies enriched for signatures C17 (**k**) and C35 (**l**). **m**, Comparing percentage of signatures C17 and C35 different clinical conditions for. Error bars in **j** and **m** indicate one standard deviation.

Different signatures are based on the high-dimensional CANVAS features that enable unbiased capture of pixel-level information. To comprehensively characterize each signature, we leveraged multiple metrics and probed the learned representation from different viewpoints.

First, we visualized the average marker intensity for each signature (Fig. 3c, see Methods). We found that marker enrichment is diverse across different signatures, including signatures enriched for a specific marker, a combination of different markers, or lack of enrichment in any markers. For marker-specific signatures, we noticed a gradient of marker intensity across similar signatures. Specifically, B cell-specific signatures (C7, C34, and C38) show a gradient of CD20 enrichment. Neutrophil-specific signatures (C49, C5, and C37) also show a gradient of MPO enrichment. Signature C0 is a T cell-enriched signature characterized by combinations of high CD3, CD4, and CD8a. Signatures C9 and C41 are examples showing no specific marker enrichment. We also found that signatures with high enrichment in specific markers usually have fewer LTMEs (Fig 3d).

Next, by deploying a set of pre-generated cell-segmentation masks, we calculated the relative cell composition in each signature (Fig. 3e, see Methods). Note that t he cell segmentation is only for interpretation purposes and not required for other downstream analysis tasks. As expected, B cell- and neutrophil-enriched signatures have a high percentage of B cells and neutrophils. The cell composition map also allows us to better characterize signatures lacking marker enrichment. Such signatures (C27, C26, C19, C12, C1, C9, C18, C41) have two distinct configurations: diverse composition or no cell enrichment. Diverse composition (C27, C26, C19, C12) has more cell types, while signatures without cell enrichment (C1, C9, C18, C41) are dominated by “None” cell types (i.e., cells that lack markers included in the panel) markers. The diverse group has lower cell density than the “none” group (Fig. 3f), suggesting that lack of marker enrichment in this group reflects a paucity of cells, while that latter group indicates lack of appropriate cell markers. These analyses demonstrate the ability of CANVAS to capture nuanced differences in marker enrichment and morphology, which is crucial for discovering and characterizing specific niches in TMEs.

Last, we investigated whether the signatures discovered by CANVAS have clinical associations. We first assessed the relative composition of signatures in patient biopsies (Fig. 3g). We grouped these samples into low and high enrichment categories for each signature, compared the clinical parameters between the two groups and identified signatures correlating with clinical variables (see Methods). Specifically, C10 has the worst prognosis but no obvious specific enrichment markers. C17 consists mostly of tumor cells but has the best prognosis. C35 is primarily associated with non-smokers while C28 is found in heavy smokers. These results raise the possibility that signatures can be used as potential biomarkers for specific clinical conditions.

### CANVAS identifies signatures representative of tissue architecture and clinical variables

We characterized LTME signatures by their intrinsic properties and found that they represent unique TMEs. To further understand their ties to tumor morphology, we mapped the LTMEs back to the original sample, and interpreted them in their tissue context. We selected C7 and C0, and C49, as example representing B cell-, T cell-, and neutrophil-enriched signatures, respectively (Fig. 3h, Supplementary Fig. 2). For each signature, we ranked patient samples by enrichment for the signature and visualized the top two samples. Notably, each signature has a distinct spatial distribution. In C7, B cell signatures form small pools and populate multiple separate regions. In C0, features are scattered and T cell TMEs have frequent interactions with B cell and tumor TMEs (Supplementary Fig. 2). We suspect that these are tertiary lymphoid structures (TLS), ectopic lymphoid organs found in tumors with germinal center-like structures (Fig. 3h, Supplementary Fig. 2). TLS are organized aggregates of immune cells that resemble secondary lymphoid organs (SLOs) anatomically, which were reported found in multiple types of cancer, including non-small cell lung cancer (NSCLC), colorectal cancer (CRC), ovarian cancer, melanoma, and others^31–35^ . TLSs are associated with favorable prognosis in several cancer types, based on the idea that in SLOs, the presence of CD4+ follicular helper T cells (TFH) and germinal center B cells indicate a robust, multiple-faceted, multicellular response. However, due to the technical challenge of identifying TLSs in clinical samples, they have not been widely used for prognostic purposes. With CANVAS, we can quickly identify possible TLS by looking at C0 enrichment in biopsies.

To further investigate spatial relationships, we selected samples enriched in a specific signature and looked at the morphology and the interactions between signatures. The T cell signature C0 generally interacts with other T cell signatures (C2, C39) as well as B cells signature C7, C34, and C38 (Supplementary Fig. 3). Compared with C38, both C7 and C34 showed higher expression of class II human major histocompatibility complex HLA-DR. This indicates C7 and C34 comprises more mature B cells with antigen presentation function. While C34 also comprises CD3, CD4 and CD8a and Foxp3 T lymphocytes markers, indicating samples with C34 might have enrichment of mature B cells with suppressive T cells, which provides a suppressive TME. B cell signature C7-enriched samples have a high concentration of B cells and similar interaction profiles as T cell-enriched samples (Supplementary Fig. 3). In neutrophil signature C49-enriched samples, other neutrophil-related signatures (C5 and C37) and tumor signatures C11, C40, C29, and C16, are present, revealing distinct morphology and interaction patterns compared with T cell- and B cell-enriched samples (Supplementary Fig. 3).

By correlating signature enrichment with clinical variables, we connected signatures to patient traits and outcomes. We found that B cell signature C7 correlates with better prognosis, confirming the correlation between B cell and good prognosis from previous studies (Fig. 3j). This might be due to the existence of HLA-DR high B cells with low Foxp3+ T regulatory cells, providing a less immune suppressive TME. Similar finding has been reported in a report examining the immune signature with breast cancer patients, showing activation of B cells that co-clustered with follicular helper T cells can mediate responses to immune checkpoint inhibitors^36^. We also found that C17 correlates with high survival (Fig. 3k,l). The biopsies with the greatest enrichment for C17 show morphologies of early-stage tumors with limited immune cell infiltration into the tumor area (Fig. 3k, Supplementary Fig. 2). In addition, we noticed that C28 correlates with heavy smoking, whereas C35 correlates with no smoking. We found that the top enriched samples for these two signatures have distinct morphologies and organizations. C28-enriched cores are typical immune-hot tumors with a mixture of T cells, B cells, monocytes, and macrophages. This was probably due to the smoking-induced carcinogen, increased somatic mutation and neoantigen that cause immunogenic response. (Fig. 3i-j). Consistent with this finding, previous studies showed that lung cancer patients who are smokers have relatively high mutational burden and neoantigen, and respond better immune checkpoint blockade ^37,38^. C35 is characterized by PanCK and HLA-DR, with limited immune infiltrates resembling an immune-cold tumor (Fig. 3l-m, Supplementary Fig. 2).

Signature C37 correlates with late-stage tumors and characterized by neutrophils with high MPO expression (Fig. 3c). Tumor associated neutrophils (TANs) have shown to correlate with angiogenesis and progression, and in mouse models, suppressive neutrophils or granulocytic myeloid derived suppressor cells (gMDSCs) that are attracted to the tumor site via CXCR1/2 signaling can suppress T cell function and promote tumor progression and resistance to EGFR/RAS/ERK pathway inhibitors^39^ . The diverse and biologically meaningful signatures identified by CANVAS demonstrated its potential to drive discovery in TME.

### Spatial organization of the TME reveals high-level tumor architectures

LTMEs can be thought of as building blocks of the TME. Thus far, we have characterized them by their marker enrichment, cell type composition, and cell density, but it remained unclear how they organize into higher level structures within the tumor. To explore this question, we embedded the LTMEs in geometric graphs to examine the properties of their interactions.

Each signature type was mapped back onto the sample to generate a signature map (Fig. 4a). We embedded the signature map into a graph by defining each LTME as a node and adding edges between nearby nodes (Fig. 4b, see Methods). To understand global interactions between different signatures, we generated an adjacency matrix based on LTME pairwise interactions across all samples (Supplementary Fig. 4, see Methods). We visualized the interactions between different signatures through a graph where larger node size and edge width represented higher LTME counts and interaction frequency (Fig. 4d). We observed that signature C9, C14, and C18, which had low marker enrichments, were the most abundant and had the most frequent interactions. They appeared to function as hubs, connecting to LTMEs with different signatures (Supplementary Fig. 5). Tumor regions, including C17, C33, and C15 also tended to interact closely. Based on the signature adjacency matrix, we performed spectral clustering and found that B cell- and neutrophil-enriched signatures formed their own clusters^40^ (Supplementary Fig. 5, see Methods). From interaction graphs, we also found that LTMEs defined by neutrophil signatures C49, C5, and C37 and B cell signatures C7, C34, and C38 had much more frequent interactions among themselves than with other signatures, confirming the spectral clustering results (Fig. 4d). In B cell- and neutrophil defined signatures, we found a hierarchical architecture wherein each signature interacts exclusively with its adjacent signature on the graph. These findings implied a topological structure, wherein the three signatures form architectures similar to concentric circles. For example, in B cell-enriched samples, this interaction goes from C7 to C34 to C38, whereas it goes from C49 to C5 to C37 in neutrophil-enriched samples (Fig. 4e-f, Supplemental Figure). The spatial adjacency of LTMEs allows us to quantitatively characterize signatures and find distinct TME architectures.

**Fig. 4:**
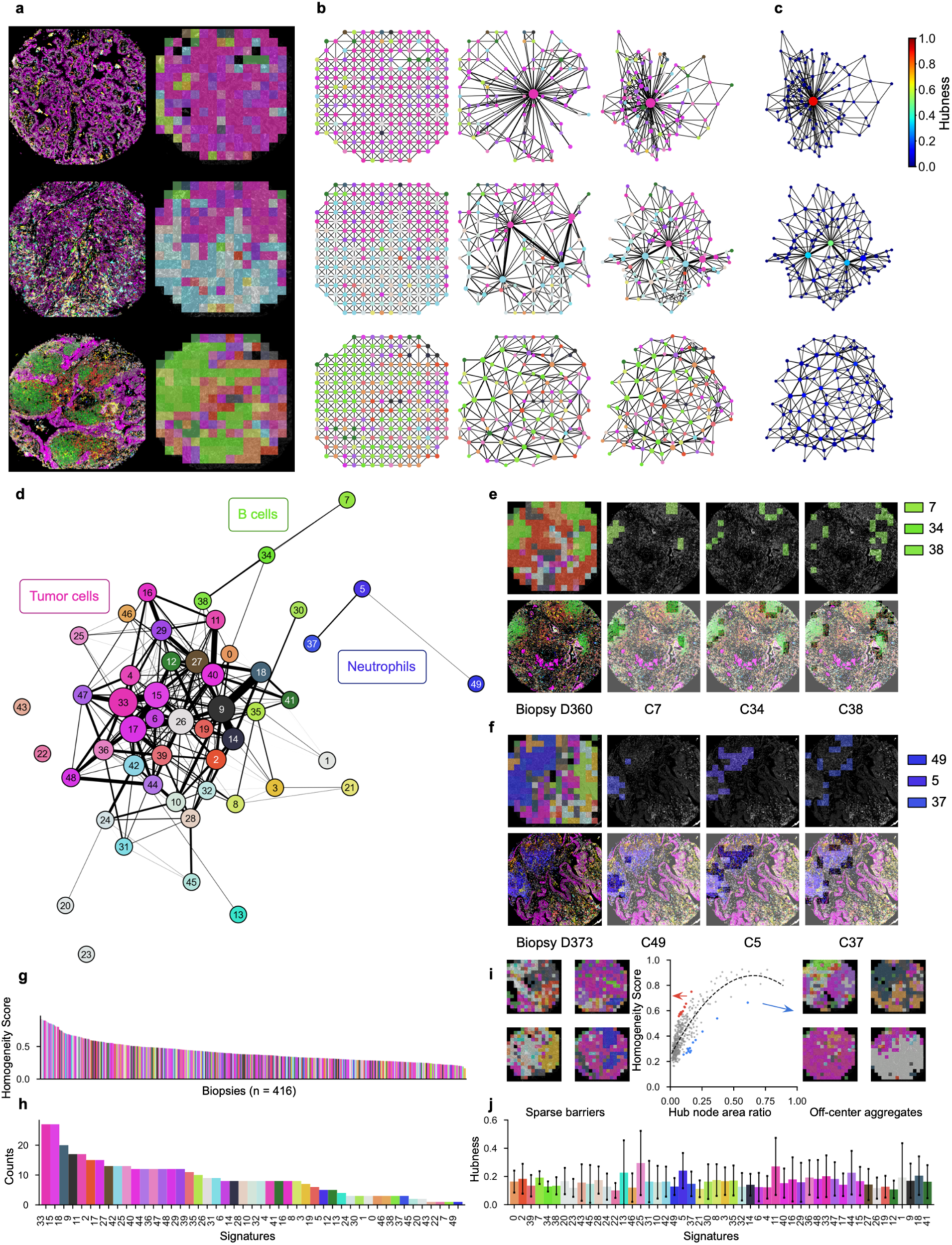
**Spatial organization of LTME signatures in tumor biopsies. a**, Original biopsy (left) and LTME signature map (right). Three separate biopsy images are colored according to the eight markers in Fig. 2c. Signature overlays are colored according to signature colors in Fig. 3a. **b**, Converting signature overlays to geometric graphs (left) with graph aggregation (middle), and layout adjustment (right). **c**, Distribution of hubness on aggregated graphs. **d**, Graph representing interactions between signatures. Node colors represent corresponding signatures. Node size denotes the number of LTMEs in each signature. Graph edges represent interactions between LTMEs from different signatures. Edge width is correlated with interaction frequencies between two signatures. Only the top 20 percent most frequent edges are shown. **e-f**, Selected biopsies enriched in B cell signatures: C7, C34, C38 (**e**), and neutrophil signatures: C9, C14, C18 (**f**). **g-h**, Homogeneity score across 416 biopsies (**g**) and the frequency of hub node signatures (**h**). **i**, Correlation between biopsy homogeneity score and hub node area (middle). Biopsies with high homogeneity and low hub node area (left). Biopsies with high hub node area and low homogeneity (right). **j**, Distribution of signature hubness. Error bars indicate one standard deviation.

### Quantification of tumor heterogeneity

One important feature of the tumor microenvironment is its heterogeneity which was shown to correlates with worse prognosis^26^ . Tumor sample heterogeneity exists at a higher level than LTMEs. To characterize how LTMEs organize and form higher order tissue architectures we need to extract the topological structures from the embedded graphs (Fig. 4b). To this end, we used the signature map, merged adjacent nodes with the same signature type, and performed snap graph aggregation (Fig. 4b, see Methods). The aggregated supernodes represent high-level structures representing different signatures and the properties of the supernodes can be used to define tissue architectures.

To characterize heterogeneity, we calculated a metric we call “hubness” for all supernodes on the aggregated graph (see Methods). We then defined the supernode with highest hubness as the hub node and its hubness as the homogeneity score (Fig 5c, see Methods). In the three selected examples from homogeneous to heterogeneous, we see a monotonous decrease in homogeneity score (Fig. 4c). We visualized the homogeneity score for all samples, colored by the hub node (Fig. 4g). We then took the hub nodes from all samples and calculated the frequency of their assigned signatures. We found that tumor signatures are the dominating hub nodes and isolated signatures like neutrophil- and B cell-enriched LTMEs are much less likely to become hubs (Fig. 4h).

**Fig. 5:**
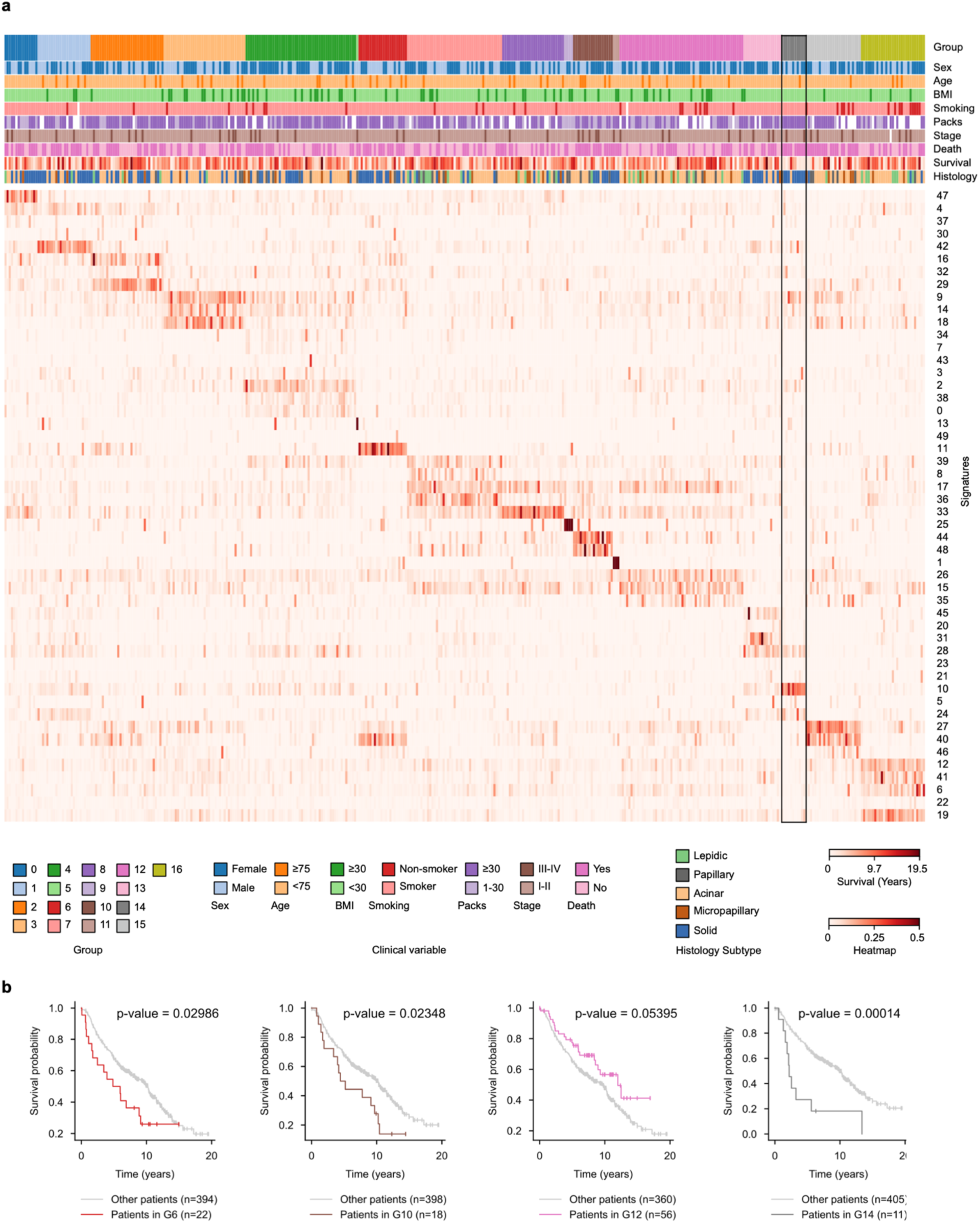
**Identifying patient groups from LTME signatures. a**, A heatmap of signature enrichment in patient samples. Patient groups are sorted by signature enrichment. Clinical variables and histology subtypes are annotated on top of the heatmap. Black box highlights the low-survival group G14. **b**, Kaplan-Meier analysis on selected patient groups.

As expected, the homogeneity score highly correlates with size of the area occupied by the hub node (Fig. 4g). However, since this score is calculated considering the topology of the sample, it captures spatial information. To demonstrate this point, we selected samples where the homogeneity score does not correlate with the hub node area percentage and identified two specific groups of morphological structures. In samples with high homogeneity score but low hub area percentage, the hub nodes must connect to as many regions as possible while minimizing the space occupied, which results in a barrier-like hub node structure that isolates different parts of the sample (Fig. 4i, left panel). In samples with low homogeneity score but high hub area percentage, the hub nodes need to maximize space occupation while minimizing connection to other regions, which results in hub nodes that are large, homogeneous regions that have limited interface with other regions (Fig. 4i, right panel).

In addition to quantifying sample heterogeneity, we can also define hubness for signatures. For each signature, we picked the supernodes with highest hubness (“top supernodes”) and used their average hubness average to represent how well each signature serves as a hub (see Methods). We calculated hubness for every signature and obtained results consistent with our observations from the signature adjacency graph (Fig. 4d, j). Specifically, more isolated signatures, including those for B cells (C7) and neutrophils (C49), have low hubness scores. By contrast, hub signature C9 and tumor-enriched signature C33, have higher hubness scores. Together, these results show that embedding signatures back to biopsies with graph analysis allows us to identify higher level topological architectures and provide a quantitative assessment of tumor heterogeneity.

### CANVAS stratifies patients into distinct subgroups

Identifying tumor subgroups can enable more customized treatments for patients. Each patient biopsy consists of LTMEs that can be categorized using the different signatures defined by CANVAS. We next explored whether the signature composition of a biopsy can be used as morphological signatures to define tumor subgroups. We performed clustering using composition of signatures as features and found 17 distinct patient groups (Fig. 5a). Different subgroups show distinct clinical outcomes (Fig. 5b). We highlight a specific group, G14, with the worst survival (Fig. 5a). Upon further investigation, we noticed that G14 also has a highly specific enrichment of the C10 signature, which we characterize in the next section. Hence, patient groups generated from CANVAS signatures could be used to guide and potentially improve diagnosis and specialized treatment.

### An immune suppressive monocytic signature is associated with low survival

In unsupervised grouping of patients, we identified a low-surviving group, G14, which is enriched in C10. We then specifically performed a survival analysis based on the enrichment of C10 and found a high correlation with poor outcome (Fig. 6a-c, Supplementary Fig. 4). C10, C28, and C45 are all enriched in heavy smokers and have similar monocytic signatures (Fig. 3g). However, unlike C28 and C45, which contain high levels of T and B cell infiltration, C10 has limited T and B lymphocytes (Fig. 6d-e).

**Fig. 6:**
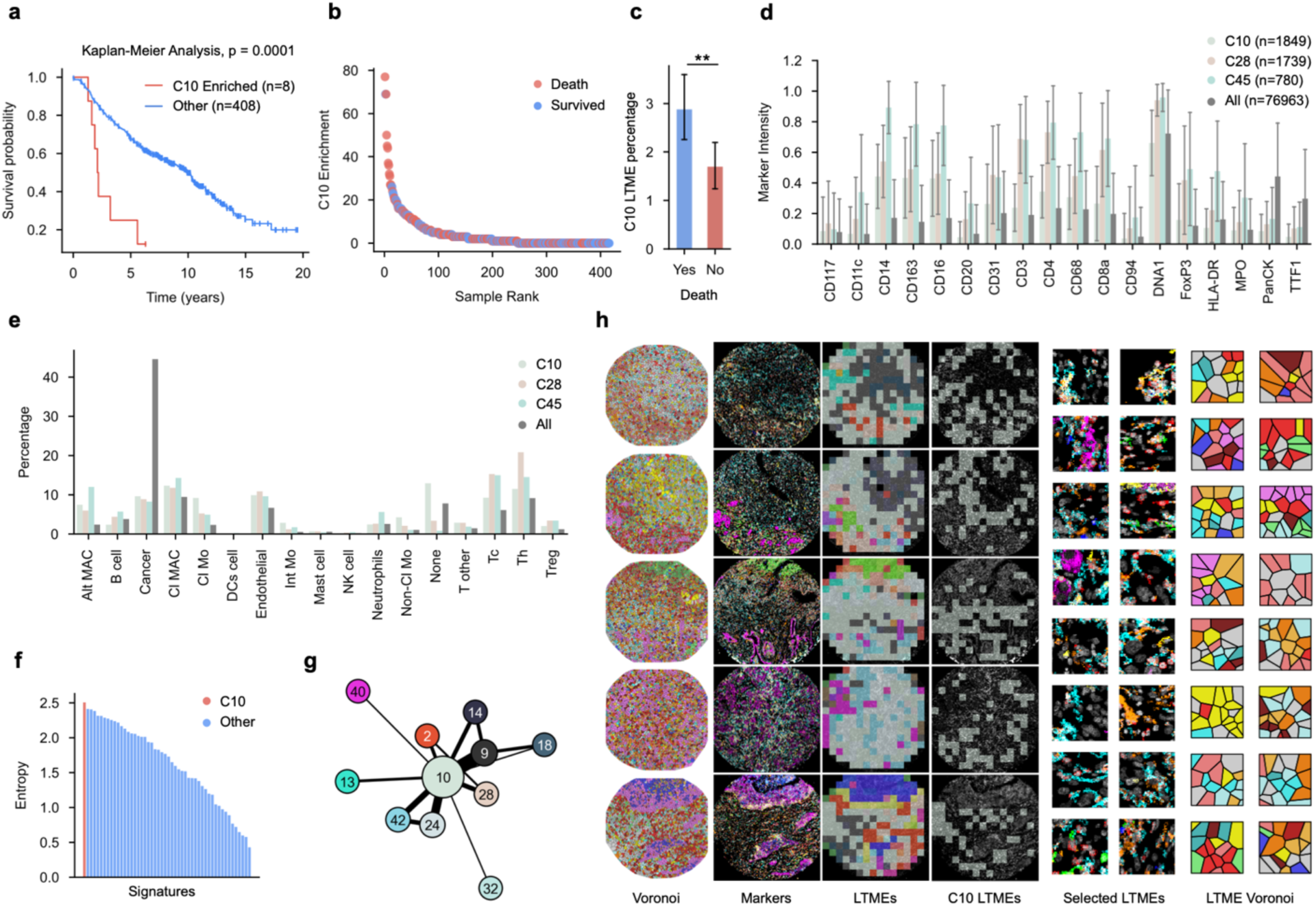
**Characterizing the monocytic signature a**, Survival analysis of C10 enriched vs other patients. **b**, Elbow plot of sample C10 enrichment colored by survival. **c**, Comparison of C10 LTME percentage between different clinical conditions. **c-d**, Marker intensity average (**c**) and cell type percentage (**d**) of C10 compared with global average. **e**, Distribution of cell type diversity across signatures. **f**, Interaction graph of signatures in C10 enriched biopsies. Node sizes and edge widths correlate with LTME counts and signature interactions frequency respectively. Only top interacting nodes are shown. **g**, Selected biopsies enriched in C10. Coloring of the cell Voronoi diagram and markers are from Fig. 3. Error bars in **c** and **d** indicate one standard deviation.

We next sought to further characterize this signature. Compared with reference markers, CD14, CD16, CD163, low HLA-DR, and low TTF and PanCK were enriched. We define this TME as a monocytic microenvironment with limited infiltration of T or B cells, and low levels (or numbers) of antigen presenting cells (APC). We also noticed that C10 has the highest cell type diversity across all signatures, which could correlate with its immune properties (Fig. 6f, see Methods). To understand C10’s interaction profile with other signatures, we examined C10-enriched biopsies (Fig. 6g). We found that C10 interacts with C9 most frequently. C9 is a common hub that connects different LTME signatures and lacks marker enrichment. C10 also interacts with C14 (Endothelial cells, no markers), C24 (CD163+ HLA-DR- M2 macrophages), C28 (immune signature with both CD4+ and CD8+ T cell infiltration), and C42 (monocyte enrichment with moderate tumor cells). These observations suggest that LTMEs in C10 are usually adjacent to connective tissue and other monocyte-enriched LTMEs, but not to tumors.

We visualized the samples enriched in C10 (Fig. 6h) and found that it is usually located at the periphery of tumors with no direct tumor interactions, which echoed the graph-based analysis. One hypothesis is that the presence of nearby tumor stimulates monocytes to migrate into these regions, which provide an immune suppressive environment that prohibits the infiltration of APCs and cytotoxic T cells. We visualized individual cells with Voronoi diagrams and analyzed the cell-to-cell interaction profile of different cell types in C10 (Supplementary Fig. 6, see Methods). Notably, most cell types in C10 had rather homogeneous interactions, compared with the concentrated interaction profile of B cell signature C7 (Supplementary Fig. 6).

### Validating the monocytic signature in an independent cohort

To test whether the C10-defined monocytic signature exists in another, independent dataset, we collected matched tumor and tumor-adjacent histological normal lung samples from 143 treatment-naive stage I lung adenocarcinoma patients at NYU. None of these patients had received any treatment for cancer prior to surgery. We performed scRNA-seq on 18 of the tumor samples and 15 of the normal samples. We also selected between one to three regions of interest from 96 of the tumor cores and analyzed them with the NanoString whole-transcriptome array (WTA) (see Methods). We then inferred cell compositions of the regions of interest with the whole transcriptome atlas generated from the scRNA-seq samples^41^ (Extended Data Fig. 1a-b).

We compared the cell composition obtained through cell type deconvolution and ranked the samples based on the cosine similarity of their cell compositions to C10 (Extended Data Fig. 1c, see Methods). Remarkably, the most similar cores were associated with a low survival rate similar to what we found in the IMC data, confirming our hypothesis that enrichment of monocytic microenvironment correlates with low survival (Extended Data Fig. 1d)

### Tumor associated monocytic upregulates ECM-related genes *in vitro*

To better understand the potential detrimental effects of this monocytic signature, we investigated its molecular mechanism and implications. Previous literature suggests that myeloid cells can have an adverse impact on patient outcomes. They could promote fibrosis in the lung under COVID settings and interact with cancer associated fibroblasts (CAFs) to exert pro-oncogenic functions in tumor^42–45^ . To examine the functionality of monocytes in the context of lung tumors, we performed single cell RNA-seq on mouse lungs and discovered that tumor associated monocytes have very different behavior from monocytes in normal lungs (Extended Data Fig. 1e-f). In particular, extracellular matrix genes are highly enriched in tumor monocytes.

To study the potential causal relationship between the TME and ECM production by myeloid cells, we performed in vitro experiments with bone marrow-derived monocytes (Extended Data Fig. 1g, see Methods). We co-cultured monocytes with or without conditioned media collected from *Kras^G12D^/Trp53^-/-^*lung tumor cultures. Remarkably, ECM expression was highly upregulated in the monocytes exposed to the tumor media (Extended Data Fig. 1h). These results provide one possible explanation for the short survival of patients with monocytic enrichment in TME. ECM produced by monocytes in TME could play a role in forming pro-oncogenic TME niches, leading to shortened survival of lung cancer patients.

## Discussion

In this work, we present CANVAS, a segmentation-free analysis framework designed for interpreting multiplexed imaging data. Representations generated by CANVAS can be used to characterize LTMEs and investigate their interconnectivity and organization in tumors. LTME composition allows partitioning of patients into subgroups with different clinical outcomes. In a large lung cancer data set, we identified a group of patients (G14) with tumors that were highly enriched in signature C10 and associated with the worst prognosis. Upon further analysis, we discovered a monocytic and immune suppressive signature in G14 and C10 and validated its adverse effects in an independent patient cohort. To understand the underlying mechanism, we used a mouse lung cancer model and found ECM-related gene upregulation in tumor-associated monocytes. Tumor-induced upregulation of ECM genes was validated *in vitro* by stimulating monocytes with tumor media. Since ECM production relates to pro-oncogenic TME niches, upregulation of ECM in monocytes might be a potential contributing factor of lower survival in monocyte enriched C10 and G14. While the C10 signature appears in both our discovery and validation cohorts, it is present in a relatively small group of patients. A larger cohort is needed to assert its association with a specific low-surviving lung cancer subgroup.

As the model is trained by using self-supervised learning, the TME representation is free of external labeling bias. However, because CANVAS is specific to its training dataset, it needs to be trained separately to be applied on different data set which could require additional computational power. This could be mitigated by performing pretraining on a large, aggregated dataset. Then the pre-trained CANVAS can be transferred and fine-tuned to new datasets with substantially less computational power. Owing to resource constraints, our in-depth application of the framework was focused on one lung cancer IMC dataset. Nevertheless, the segmentation-free nature of CANVAS makes it feasible in a wide range of imaging analysis under different conditions, including multiplexed immunofluorescence imaging, imaging mass spectrometry, and spatial transcriptomics. Even though CANVAS is trained in a stochastic manner, the LTME features are robust and consistent across different runs (Supplementary Fig. 6-7). CANVAS is orthogonal to existing cell-segmentation-based analysis. It is possible to combine segmentation-based and segmentation-free methods to provide a more comprehensive understanding of imaging data. In addition, we are aware that masked image modeling is one of the many self-supervised learning algorithms to learn image representations. It is possible to implement other techniques like contrastive learning for the same purpose. However, the resulting embeddings could differ since each self-supervised learning implements a different learning objective. We will leave that for future studies to explore.

The morphology and organization of cells are essential components that contribute to the complexity of TMEs. The identification and characterization of distinct TMEs demonstrate the value of the CANVAS framework in understanding the heterogeneity of TMEs. In addition, CANVAS can independently generate novel clinical and biological hypotheses that can be validated through both external dataset and *in vitro* experiments. We expect future work to leverage pixel-level approaches as the analysis framework to capture the full complexity of high-dimensional spatial molecular measurements of tissues.

## Glossaries

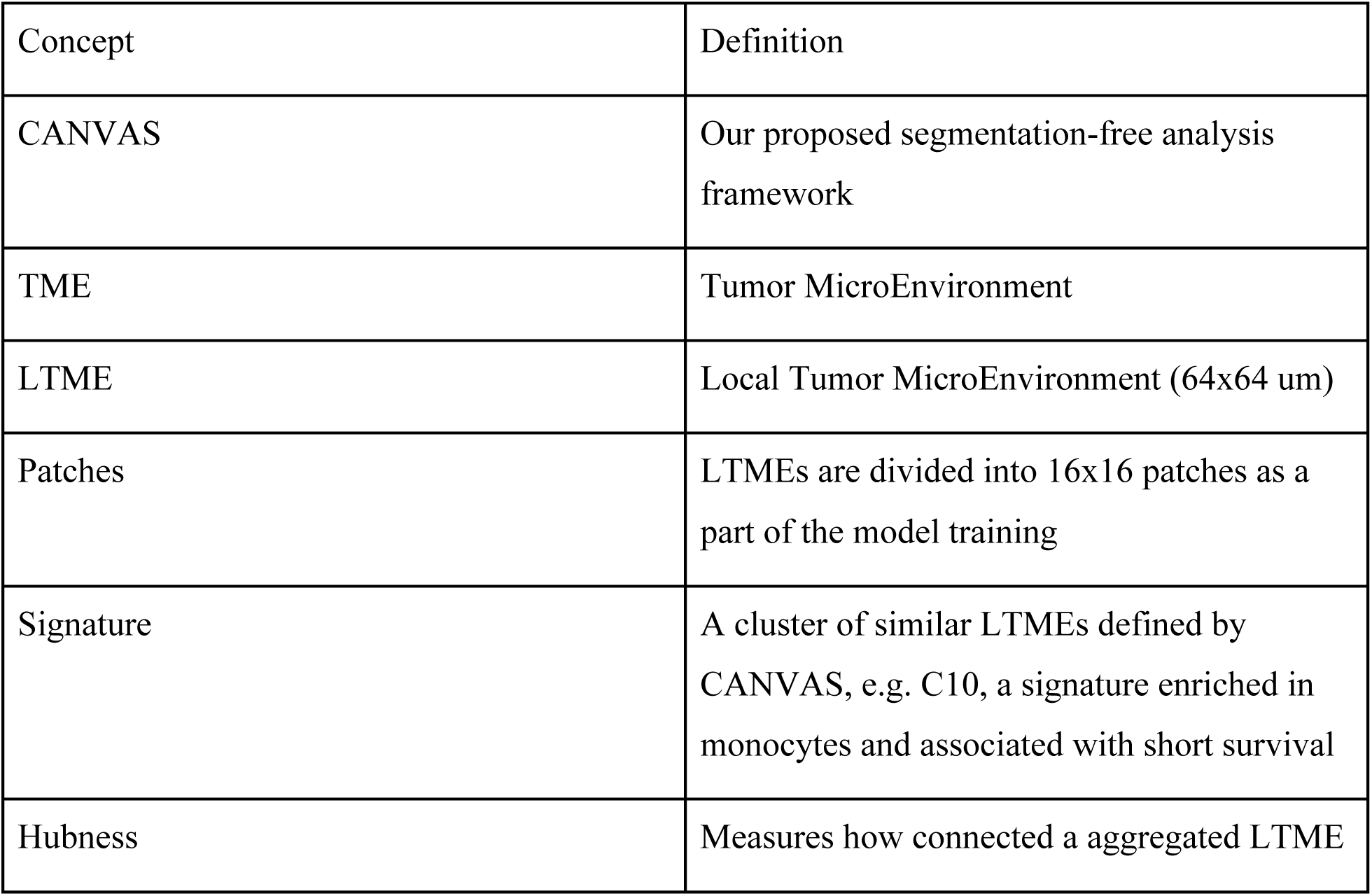
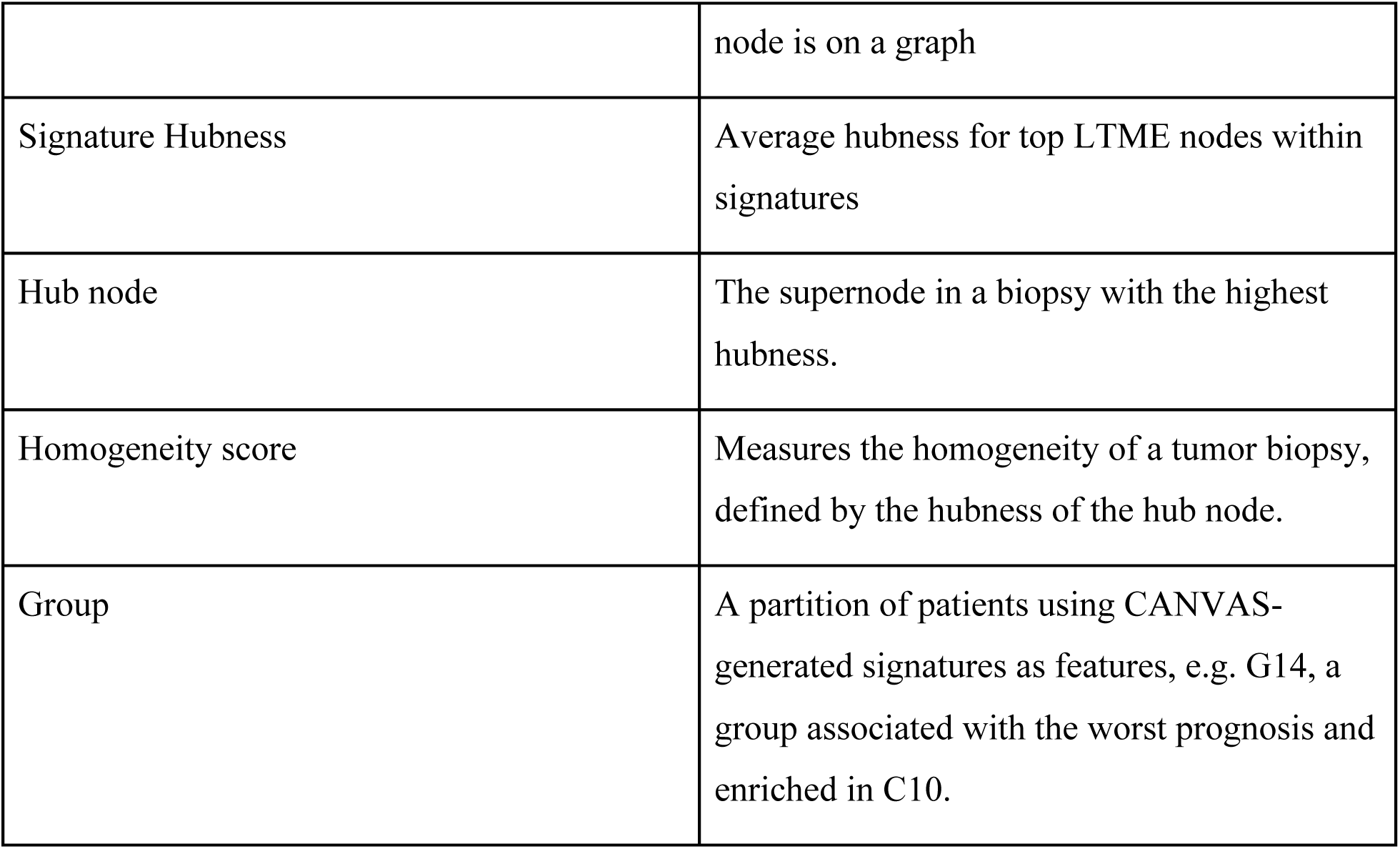

| Concept | Definition |
| --- | --- |
| CANVAS | Our proposed segmentation-free analysis framework |
| TME | Tumor MicroEnvironment |
| LTME | Local Tumor MicroEnvironment (64x64 um) |
| Patches | LTMEs are divided into 16x16 patches as a part of the model training |
| Signature | A cluster of similar LTMEs defined by CANVAS, e.g. C10, a signature enriched in monocytes and associated with short survival |
| Hubness | Measures how connected a aggregated LTME |
|  | node is on a graph |
| Signature Hubness | Average hubness for top LTME nodes within signatures |
| Hub node | The supernode in a biopsy with the highest hubness. |
| Homogeneity score | Measures the homogeneity of a tumor biopsy, defined by the hubness of the hub node. |
| Group | A partition of patients using CANVAS-generated signatures as features, e.g. G14, a group associated with the worst prognosis and enriched in C10. |

## Methods

### Preprocessing of IMC image data

The LUAD IMC data is obtained from the zenodo public repository (https://zenodo.org/record/7760826). The repository contains samples and clinical information from 416 patients with lung adenocarcinoma. For each patient, we downloaded the IMC image, preprocessed cell segmentations and clinical data. Each image file is first converted to numpy tensors. Content masks are generated to avoid feeding empty regions to model and increase learning efficiency. To generate a mask, we first averaged across the channel dimension and downsampled the original image on x and y dimension with a factor of 16 to speed up processing. We then generated a mask by adjusting the pixel value range to [0, 1] and selecting regions with values larger than a threshold of 0.2. To generate tiles for training, we selected a square tile of 64 by 64 micron. The IMC data is acquired at a resolution of 1 pixel per micron. 64-micron tile is selected due to cell size and the masking strategy in masked imaging modeling. In masked imaging modeling, the image is separated to 16 by 16 grids and 256 patches in total. During training, the patches are masked randomly. In a 64 by 64-micron image, a patch has dimension 4 by 4 micron, which is roughly less than a quarter of the area of a cell with 10-micron diameters. By choosing a 64-micron tile, we implicitly enforce the model to encode information at cellular level. For every patient sample, we separated it into 64-micron image tiles. After tiling, each image is a neighborhood containing around 20 cells. We then filtered the tiles by averaging the mask value within the tile and removed the tiles with average intensity less than the same threshold of 0.2.

### Model architecture

CANVAS is an asymmetric autoencoder consisting of a standard ViT (Vision Transformer) encoder and a lightweight ViT decoder. The overall architecture is similar to the model described in Masked Autoencoder^30^ . The encoder is a standard ViT-Large with 24 transformer blocks, 1024 hidden size and 16 multi-head attention^29^ .

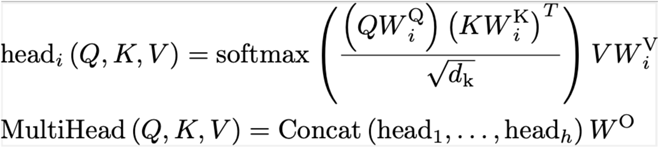

The decoder is a lightweight ViT with 8 blocks and 512 hidden size. A 2D sin/cos positional embedding is added to the input to encode spatial position of patches. The input dimensions are changed to 18 to accommodate the 18-plexed IMC image data.

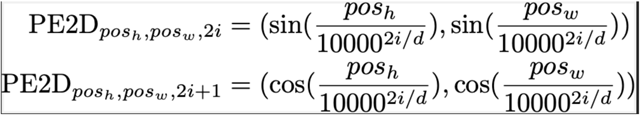

### Model training

Since CANVAS is a framework geared towards data exploration and discovery with self-supervised learning encoding and unsupervised clustering, data leakage does not apply. Therefore, the model is trained on all available tiles. During training, the tiles size used is different from inference time because the training tile size needs to leave room for data augmentations. Thus, we used a 128-by-128-micron image as input. The augmentation will reduce the image size by half on average. Then the 128-by-128-micron image will be reduced to roughly to a 64-by-64-micron (64-by-64-pixel) region. This tile is then resized to 224 by 224 pixels and split into 256 14- by-14-pixel patches. In the encoder, each input image patch is encoded by a convolutional neural network (CNN). The positional encoding is directly added to the patch embeddings. Then, we mask out 192 (75%) patches and keep them as reconstruction targets for decoders. The rest of the patches are reorganized into a sequence of features and fed into the transformer encoder. The encoder-processed patch embeddings are then combined with masking tokens and restored to their original position on the image. Then an additional positional embedding is added. The output is then fed into the lightweight transformer decoder to generate the final predicted patches. The loss is calculated by comparing predicted masked patches to the original masked image patches with an MSE loss. The model is pre-trained on tile from all 416 samples for 2000 epochs on a GPU workstation equipped with 4 NVIDIA Tesla V100 GPUs for 48 hours.

### Data normalization and augmentation

Marker intensity across different imaging channels varies due to biological variations and imaging artifacts. CANVAS relies on MSE reconstruction loss to perform representation learning. We performed channel-wise normalization by sampling 10000 random tiles and calculated the channel-wise mean and standard deviation for the whole dataset. During training, inputs to the model are first normalized using the dataset statistics. To increase robustness of the model, we added various data augmentation strategies that reflect biological and technical variations of the source data. The augmentation step includes random rotations from zero to 180-degree, horizontal flip, random cropping to 50-100% of the original scale and channel intensity augmentation. The channel intensity augmentation multiplies each channel with a random variable sampled from gaussian distributions. The normalization and augmentation of each channel is implemented as below:

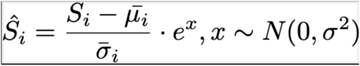

Where 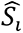 is the data pre-normalization and augmentation. 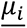 and 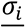 denote the pre-calculated dataset mean and standard deviation. 𝑒*^x^* is the augmentation term where 𝑥 follows a gaussian distribution. In our experiment, setting sigma to 0.3 achieves a good effect.

### Embedding visualization and clustering

Tumor biopsies from 416 patients are separated into 76963 individual LTME tiles. The trained CANVAS model is then used to generate embeddings on all tiles without masking. The model produces a 14 by 14 by 1024 channel tensor for each tile. We reduced the tensor’s two spatial dimensions by averaging across the first two axes, resulting in a 1024-dimensional vector for each tile.

To visualize the embeddings, we performed dimensionality reduction on the embeddings using UMAP. Then we superimposed the original tile image onto the location of its corresponding embedding in the UMAP space. For each superimposed image, we visualized 8 representative channels (CD20, CD3, CD14, CD68, Nuclei, MPO, HLA-DR, PanCK) out of all 18 channels with different colors.

The embeddings are then clustered with the K-means clustering. We tested numbers from 3 to 130 with the elbow method to optimize the number of centroids. However, no sharp elbow point was identified. Thus, we selected 50 centroids to capture enough between-group diversity while not over fragmenting the biological groups.

To generate a heatmap of signatures by markers, we first performed a min-max scaling on the original IMC image. 2 and 98 percentile of pixel values across sample and channels are selected as the min and max values to remove extreme values and increase dynamic range. Then we selected LTME tiles that belongs to each signature and averaged marker intensities across tiles.

### Incorporating cell segmentation and clinical variable

Cell segmentation data is obtained from the original zenodo data repository (https://zenodo.org/record/7760826). We mapped cells onto each tumor biopsy and further to each LTME tile. Similar to the marker enrichment analysis. We counted the number of cells for each cell type across all tiles sharing the same signature and plotted cell type percentages against signature.

To find out how signatures correlate with clinical variables, we need to find a way to connect signatures to samples. For each signature, we counted the number of corresponding LTME tiles in each sample and divided it by the total number of tiles to find the tile ratio, or relative enrichment of the tile in each sample. Then we group all the LTME tiles by the clinical variable its original sample has. For instance, if we are looking at smoking status, a LTME tile is labeled smoking if the tumor biopsy that contains this tile has the smoking label. Then we looked at the distribution of enrichment and assessed the difference between two groups. For each pair of groups (e.g., Smoker vs Non-smoker), Mann-Whitney U tests were conducted between two groups to compare their enrichment distributions. False discovery rate (FDR) correction was applied to the p-values within each signature using the Benjamini-Hochberg procedure. The adjusted negative base-10 logarithm of the FDR adjusted p-values was computed for each test.

### Spectral clustering on global signature adjacency matrix

A global adjacency matrix 𝐴 ∈ 𝑅^$×$^ of all signatures is created by aggregating pair-wise edges between different LTMEs in signature maps across all samples, where 𝑐 is the number of signatures. Then, we define diagonal matrix D whose diagonal elements are the weighted degree of each signature vertex. The weighted degree can be computed by summing over the rows in the adjacency matrix for each signature vertex:

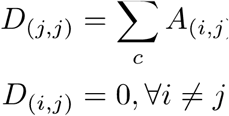

Then the graph Laplacian L can be computed as the difference between weighed degree matrix D and adjacency matrix A:

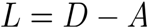

Then, given the desired number of clusters n, we find the top n most important eigenvectors

{𝑥_&_, … 𝑥_’_} ranked by corresponding eigenvalues. These eigenvectors can be stack to a matrix 𝐸^n×$^ and normalized for each signature. Then, each column in the normalized 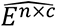 is used a feature for Kmeans clustering using 𝑛 as number of centroids and each signature will be assigned to a centroid.

### Generation and aggregation of LTME interaction graphs

To generate the interaction map between different LTMEs, we first mapped each LTME to their original position on the tumor biopsy. Each LTME is defined as a node n. Then we connected nearby nodes with edges, which resulted in a set of tumor geometric graphs. We then aggregated all the edges across samples and calculated the pairwise frequency between nodes with different signatures. This contact frequency matrix was then converted to a contact topological graph with node side denoting the number of LTMEs in a signature and width of the edges between signatures denoting the number of their pairwise interactions. For a clean visualization, we only visualized edges with numbers of interactions in the top 20 percentile.

To investigate the higher-level topological structures of tumor tissue architecture, we performed graph aggregation to reduce the number of nodes in each graph. The aggregation algorithm iteratively merges adjacent nodes with the same signature. Formally, let all nodes on the graph as a set 𝑁 and a subset of all nodes with signature n as 𝑆_’_. We define the merge operation as ⊕ and adjacent node as 𝑢 − 𝑣, one supernode generated from one pass of the algorithm can be described as:

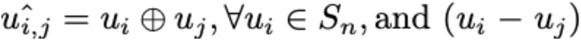

After aggregation, the edge set of the supernode 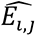 is the union of the edge set of the precursor nodes.

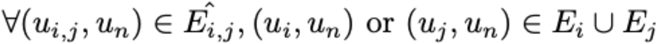

This algorithm is run iteratively until there are no nodes left with adjacent nodes from the same signature. After the merge operation, the remaining structure could include repetitive subgraphs. We additionally performed SNAP summarization to identify and purge isomorphic structures in the graph for more succinct representations of the topology^46^ .

### Hubness and homogeneity score

A single LTME node only connects to its nearby nodes. After aggregation, the supernodes retained all the edges from their precursors, allowing them to have many connections. We expect supernodes to have different connection profiles. Some nodes can be isolated while the other nodes act as hubs connecting other nodes. We defined a concept we call hubness to quantify how connected each supernode is. Hubness of a node is defined as the inbetweeness of a node on a graph. Inbetweeness is a common graph centrality measurement that can be defined as follows:

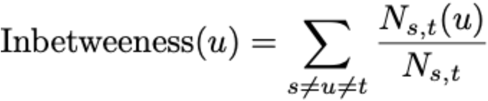

Where 𝑁*_s,t_* is the total number of shortest paths from node s to node t and 𝑁*_s,t_*(𝑢) is the number of paths from 𝑠 to 𝑡 that also pass through 𝑢.

The hubness score can be defined for signatures. Within each signature, we calculate the hubness of each LTME supernode. The top 10 percent of the supernodes with high hubness are selected. The average hubness of the top supernodes is defined as the hubness of the signature.

Since hubness captures morphological information, we used hubness of LTME supernodes to define a homogeneity score for the tumor tissue. We define the homogeneity score of a tumor biopsy as the maximum hubness score of all the LTME supernodes in the tumor. If the homogeneity score is high, this usually signifies a dominant signature that occupies the whole tumor biopsy which correlates with higher homogeneity.

### Generating cellular tessellation and graph representations

Cell segmentations are mapped onto tissue biopsies. The voronoi diagram is generated by calculating the Delaunay Triangulation and coloring each region by cell type. The cellular interaction matrix is generated by constructing a Gabriel Graph which is more stringent and avoids connecting remote interactions. The graph aggregation with SNAP summary is performed in the same way as described in the LTME graph analysis section.

### LUAD Patient Population and Tissue Microarray construction

Thirty-one IASLC grade 3 Stage I adenocarcinomas with at least five years of follow-up were chosen from the NYU Langone Health Thoracic Surgery Archives. Representative formalin fixed paraffin embedded tumor and uninvolved lung at least 5 cm from the tumor were used to construct a TMA by the Center for Biospecimen Research and Development of NYU Langone Health. A pathologist (AM) annotated each case and region that was used to generate 2 mm diameter cores for each TMA block. Each block was sectioned and HE stained for review by the pathologist and scientists annotating regions of interest for DSP.

### NanoString GeoMX Digital Spatial Profiling

Each TMA block was sectioned with precautions to avoid introducing RNAses and decrease the time from section to experiment. Briefly, sections were recovered in RNAse free water and stored at 4° C in a sealed box to avoid humidity and condensation. All incubations and washes were carried out following NanoString® standard of practice. Briefly, after baking for 1-2 hrs at 60° C sections were deparaffinized and antigen retrieved (20 minutes at 100°C) with EDTA based solution (ER2 Leica Cat. AR9640) for RNA assays and citrate based solution (ER1 Leica Cat. AR9961) for protein assays on a Leica Bond Rx Autostainer. For RNA assays protease K was used for retrieval (1 µg/mL for 15 min). After antigen retrieval slides were moved to PBS for probe or antibody incubation overnight. For whole transcriptome analysis slides were incubated with WTA probe mix consisting of ∼18000 annotated probes with photocleavable barcodes (200 ml) covered with a Hybrislip (Grace biolabs) in an HybEZ™ oven at 37°C followed by two stringent washes (2XSSC, Formamide 50% v/v, at 37°C) and two 2XSSC (room temperature) washes. The slides were then immunofluorescently labeled with RNA compatible solid tumor morphology markers (PanCK, CD45, SYTO13, NanoString® cat.

121300310). The slides were then loaded onto the instrument for region of interest selection. Tag acquisition was performed after fluorescent scanning of each slide at 20X magnification (0.4 mm/px). Three regions of each TMA core were selected for segmentation into panCK(+) and panCK(-) segments that each contained at least 100 cells for RNA assays or 30 cells for protein assays. Each segment was processed individually, and UV-released barcodes were collected into 96-well microplates for downstream Illumina sequencing.

The sample collection plate is sealed and stored at -20 deg Celsius. After thawing, the plates are sealed with AeraSeal film (Excel Scientific, CA) and the aspirates are dried at 65 deg C in a thermal cycler with an open top. Once dried, the wells of the collection plate are rehydrated with 10ul of DEPC water, mixed and spun. 4ul each of the DSP aspirate and primer (GeoMx SeqCode Primer plate) are added along with 2ul of the the 5X PCR Master mix to a new PCR plate and mixed. The sealed plate is transferred to a (Bio-rad C1000 thermocycler (Bio-rad, CA) and Illumina flowcell compatible libraries are constructed by 18 cycles of PCR amplification according to instructions.

4ul each of the resultant libraries (from each well of the PCR plate) are pooled into a 1.5 ml tube and cleaned using AMPure XP beads (Beckman Coulter Life Sciences, IA). The quality of the pooled libraries is assessed on tapestation, where the amplicon size is 162 bp and no residual primers are detected. Post quantification of them on a qPCR, the libraries are sequenced on a NovaSeq6000 SP100 flowcell (Illumina Inc., CA) as paired end reads with 5% PhiX spike-in.

The resultant sequencing reads are demultiplexed using the barcode information provided and fastq files are generated. The GeoMx NGS pipeline (DND software from nanoString) is then used to generate digital count conversion (DCC) files. The zipped DCC folder is then transferred to digital spatial profiler (DSP) and downstream data analysis performed.

### BayesPrism Deconvolution

We generated the cell type annotations for our scRNA-seq dataset using reference annotations from a previous study of lung adenocarcinomas^47^ . Brain metastasis samples were filtered out from the reference dataset. Using SingleR^48^, cluster profiles were aggregated, and each of the 86 clusters at resolution 3 were annotated with a cell type and cell subtype. Seurat anchor-based integration for merging datasets was applied on 15 normal samples (51,248 nuclei) and 18 tumor samples (61,210 nuclei) for biological and technical batch correction between patients^49^ . This method identifies pairwise correspondences between cell pairs across datasets to transform them into a shared space. Two dimensionality reduction steps were performed. The 2,000 most variable genes were selected for canonical correlation analysis (CCA). The first 30 dimensions were then defined as the integration anchors. Cell gene expression was quantified using Seurat’s AddModuleScore function. We filtered out ribosomal and mitochondrial genes, genes on the sex chromosomes, as well as genes expressed in less than 10 cells, leaving us with a total count of 13,972 genes.

Using the annotated scRNA-seq from our NYU tumor samples as single-cell reference, we applied BayesPrism on each region of interest (ROI)’ whole transcriptome atlas and inferred their cell-type composition^41^ . BayesPrism’s parameter key was set to NULL indicating the absence of malignant cells in the reference dataset and the equal contribution of all cell types. The final cell type proportions were estimated with Gibbs theta values.

### Transferring LTME signature with cell type composition

To find the regions of interest that are mainly composed of Cluster 10, we compared the cell-type composition of each ROI and that of Cluster 10 using cosine similarity rank. For the cell annotations to match, we adjusted the cell annotations for both the Cluster 10 cell types and the ROI’s cell types from BayesPrism. We removed the AT1, AT2, Ciliated, Club, COL14A1-matrix- FBs, COL13A1-matrix-FBs, myofibroblasts, pericytes and CD207CD1a-LCs cells from our ROI’s cell annotations, and the None and Neutrophils category from the Cluster 10 composition. For the Cluster 10 cell types, we grouped Alt MAC and Cl MAC under macrophages, Cl Mo, Non-Cl Mo and Int Mo under monocytes, T other, Tc, Th, and Treg under T cell. For the ROIs’ cell types, we grouped CD1c-DCs and Activated-DCs under DCs cell, Tip-like-ECs, Stalk-like-ECs, and Lymphatic-ECs under endothelial cells, and renamed Alveolar-Mac as macrophages, Tumor as cancer cells, T-lymphocytes as T cell and B-lymphocytes as B cell. We normalized Cluster 10’s cell composition and all ROIs’ cell type proportions so that they each amount to 1. We computed cosine similarity scores between each ROI and Cluster 10’s cell composition and ranked the ROIs from the highest score to the lowest. We selected all ROIs with a cosine similarity score higher than 0.625 and labeled them as Cluster 10-matched ROIs. For progression-free survival (PFS) analysis of Cluster 10-matched ROIs, we stratified the ROIs into Cluster 10-matched ROIs and other ROIs and performed progression-free survival analysis using PFS log-rank test.

### Single-cell RNA Sequencing data processing

Filtered cell matrices of naïve and tumor samples from CellRanger were imported into Scanpy for preprocessing and analysis^50^ . Cells having less than 200 or more than 9000 genes detected, or more than 20% of total UMIs stemming from mitochondrial transcripts were removed from further consideration. Raw counts were normalized by dividing the count by the total counts per cell and then log-transformed with a pseudocount of 1. Top 3000 highly variable genes were selected for PCA calculation. UMAP projection and Leiden clustering were based on top 40 components. Batch corrections were performed using Harmony with default parameters^51^ . Marker genes of each cell clusters were identified using *rank_genes_groups* function available in Scanpy. Cell type annotations were conducted manually based on canonical cell type markers and literature search.

### Monocyte cell culture and RT-PCR analysis

Bone marrow aspirates were harvested from mouse tibias and femurs by flushing them with ice-cold complete RPMI (containing 10% FBS, 2mM Glutamine, and 100U Penicillin/Streptomycin). This was followed by red blood cell lysis and filtration using a 70µM cell strainer. 8.5 x 10^5 single bone marrow progenitor cells were plated in monocyte/macrophage media (complete RPMI media supplemented with 20ng/mL M-CSF) on a 10cm petri dish and allowed them to grow in culture for 4 days. On day 5, the monocyte/macrophage media were replaced with tumor conditioned media collected from KP tumor cell cultures (*Kras^G12D^/P53^flfl^*, Clone C) for the time points as indicated.

Total RNA was extracted using the RNeasy plus Kit (Qiagen) and converted to cDNA using qScriptTM_cDNA_SuperMix (Quanta Biosciences). Real time qPCR was performed with primers and iQTM SYBER Green master mix on a QuantStudio 5 (Applied biosystems) by a standardized protocol. Primers used for specific genes are as follows:

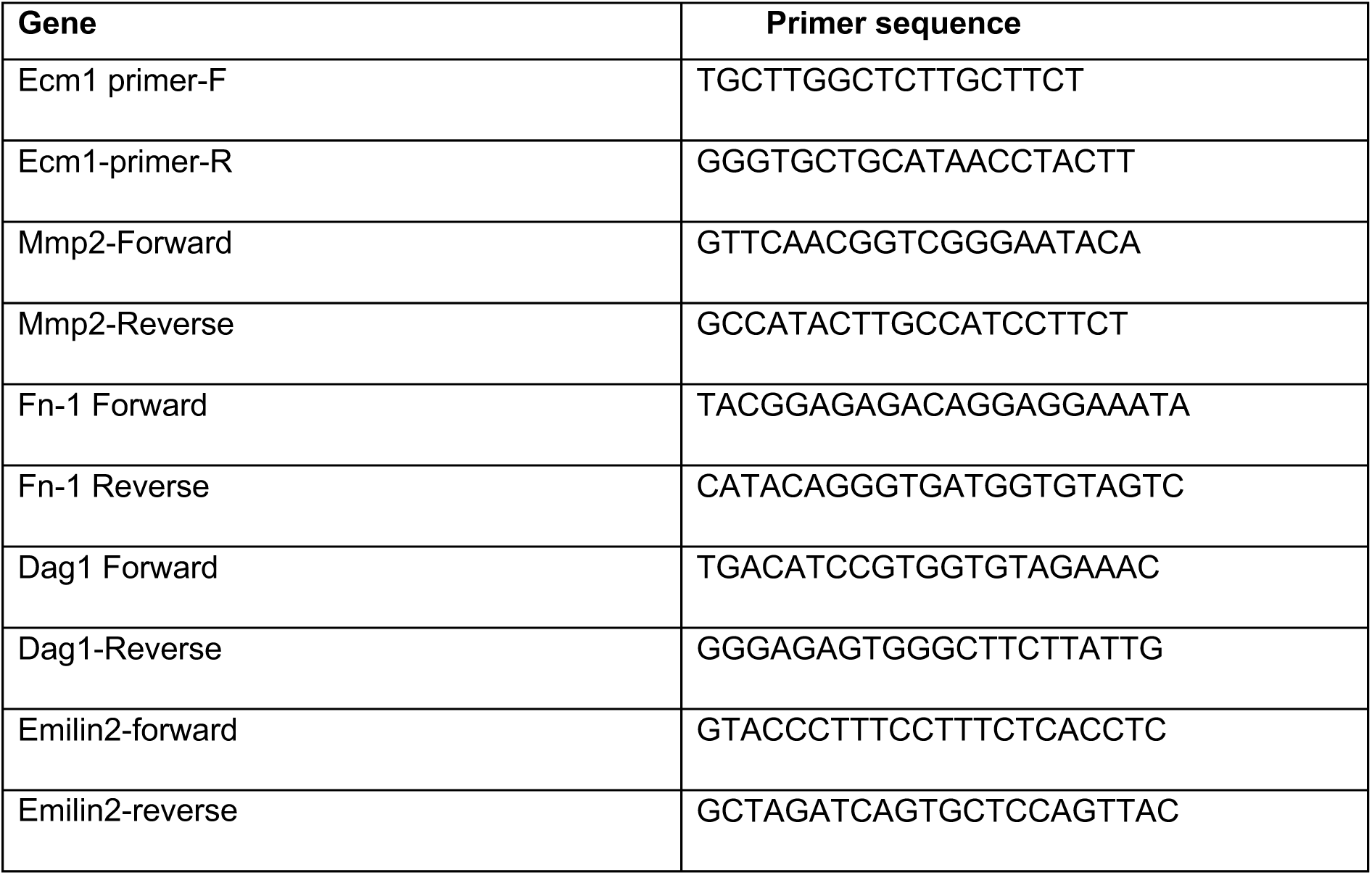

## Data availability

The IMC images, cell segmentation and patient metadata used in the study were public data from zenodo with access link listed in the Methods section. The single cell monocyte and qPCR data will be uploaded to GEO. The NanoString GeoMx data will be uploaded to zenodo data portal.

## Code availability

The code used for CANVAS training and downstream analysis is available at https://github.com/tanjimin/CANVAS.

## Acknowledgements

J.T. and D.F are supported by P01AG051449. A.T. is supported by the Cancer Center Support Grant P30CA016087 at the Laura and Isaac Perlmutter Cancer Center. Experimental Pathology [RRID:SCR_017928] is partly funded by NIH/NCI 5 P30CA16087. This work has used computing resources at the NYU School of Medicine High Performance Computing (HPC) Facility. We thank the Adriana Heguy and Genome Technology Center (GTC) for the support on expert library preparation and sequencing. We thank Harvey I. Pass for the assistance on lung tumor data collection.

## Author Contributions

J.T. and D.F conceived the project. J.T, A.T. and D.F designed the experiments. J.T designed and implemented the model and framework and performed downstream analysis with inputs from B.X., Y.B., W.L. and J.M.W. S.R., V.M. and C.L. performed GeoMx WTA experiments. H.L. and N.M. analyzed the GeoMx WTA data. J.D., K.W., Y.B., J.T., A.L.M., B.G.N., A.T., and D.F interpreted results. Y.B. performed mouse single cell experiment. Y.B. and Y.L. designed and performed the monocyte experiment. Y.H. performed mouse single cell analysis. J.T. prepared figures with input from H.L. B.X. A.T. and D.F. J.T, J.D., B.G.N., A.T. and D.F. wrote the manuscript with inputs from all authors.

## Inclusion & Ethics

We confirm that all authors are accredited according to their contribution to this work. Analysis in this study is based on publicly available datasets with no bias in selection related to subject diversity.

---

**Extended Data Fig. 1:**
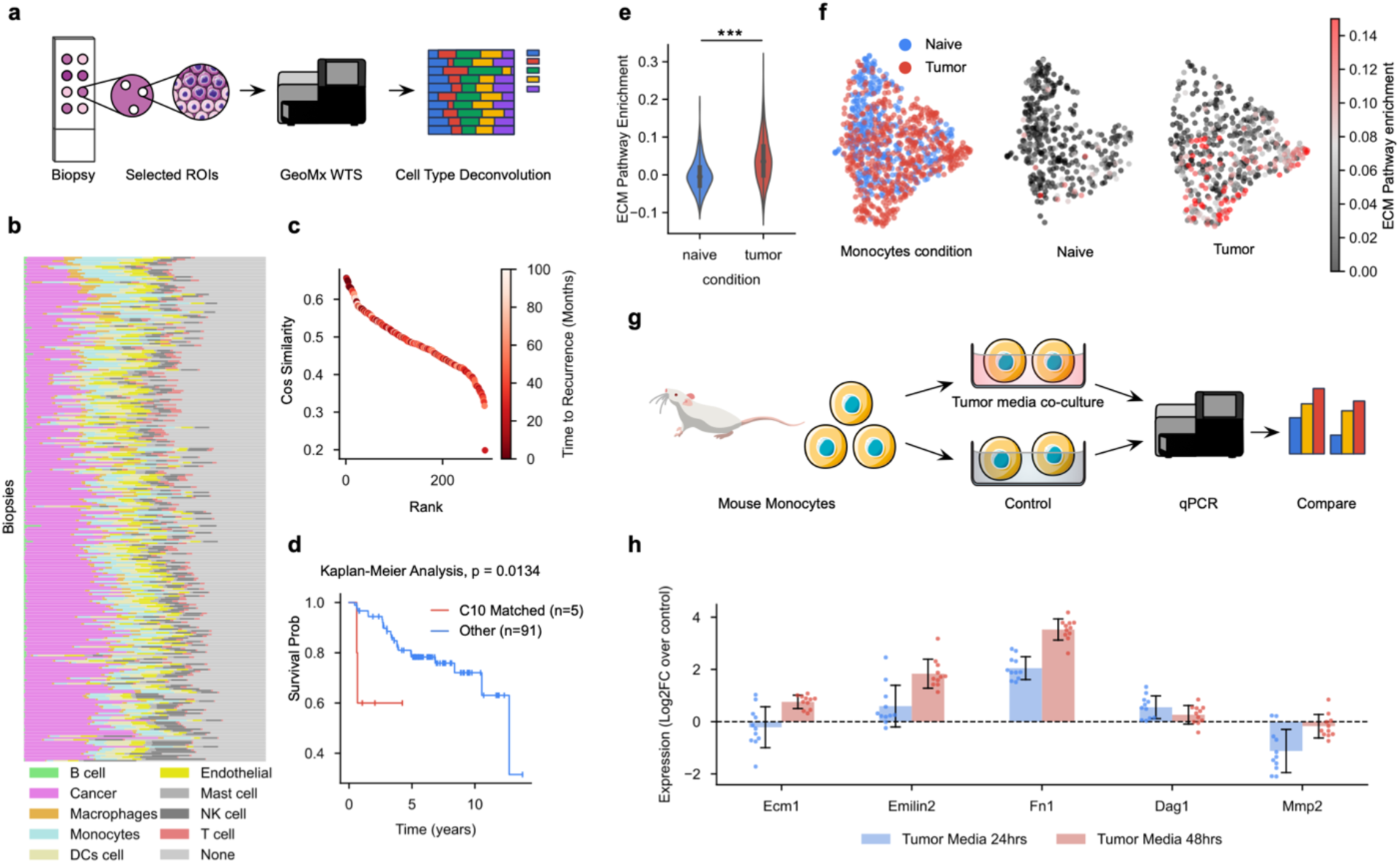
**Validation of the monocytic signature. a**, Schematic of WTA data processing pipeline. **b**, Cell type composition across biopsies. **c**, Elbow plot of biopsies ranked by cosine similarity to signature C10, colored by survival. **d**, Survival analysis of C10-similar patients vs other patients. **e**, Cell level enrichment score comparison between naïve lung and tumors. Error bars in violin plots indicate minimum, mean, and maximum values within each group. **f**, Single cell analysis on tumor and normal monocytes. Distribution of monocytes in tumor and naive mouse lung (left). Extracellular Matrix (ECM) pathway gene expression in naive (middle) and tumor (right) lung, colored by enrichment score. **g**, Schematic of in vitro monocyte co-culture experiments. **h**, Log2 fold-change of ECM-related gene expression of the treatment group (24 and 48 hours) compared with the control group.

**Supplementary Fig. 1:**
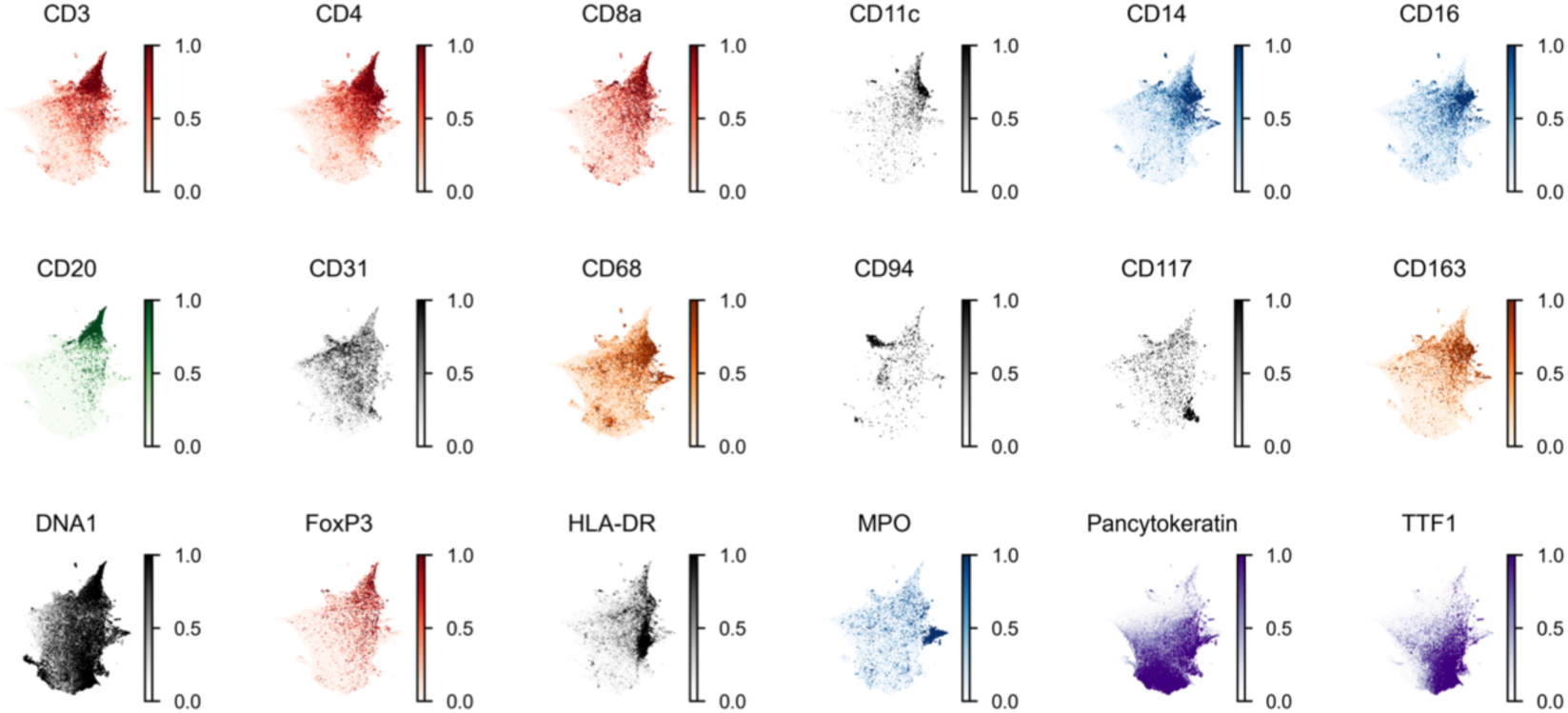
Distribution of markers on CANVAS embedding. The UMAPs of CANVAS LTME embeddings are colored by average intensity of the 18 markers shown.

**Supplementary Fig. 2:**
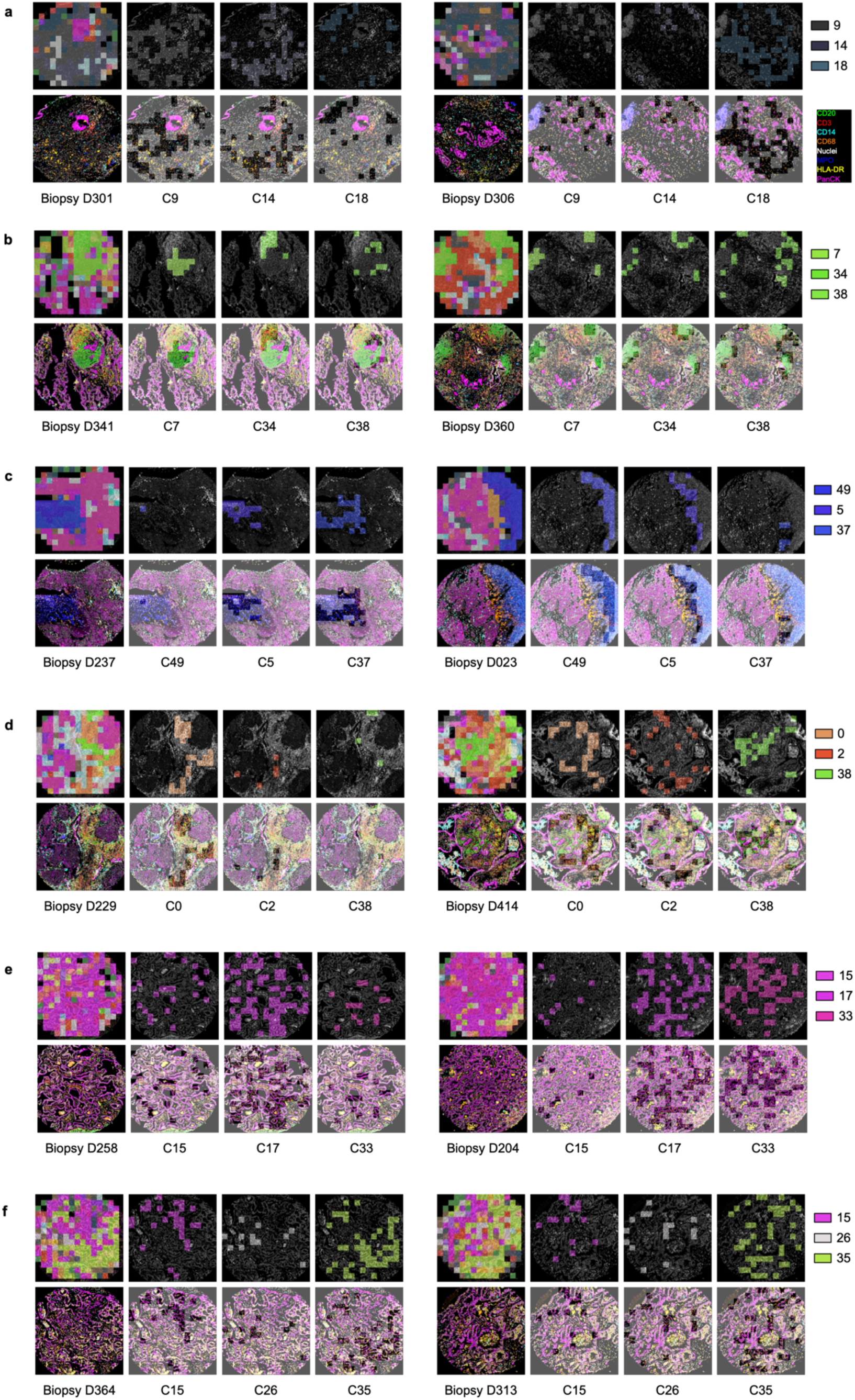
**Selected biopsies with high signature enrichment. a-e**, In the first column, LTME are denoted by their signature color. The bottom image is rendered using the default palette. Distribution of specific LTME signatures is shown from the second to the last columns. Top row: LTMEs are shown by their signature colors. The image is rendered in grey scale by averaging all the color channels. Bottom row: images are overexposed except for selected LTMEs. We show signatures that we define as no marker regions (C18, **a**), B cell-enriched (C7, **b**), neutrophil-enriched (C49, **c**), T cell-enriched (C0, **d**), good prognosis (C0, **e**), and non-smoking (C35, **f**).

**Supplementary Fig. 3:**
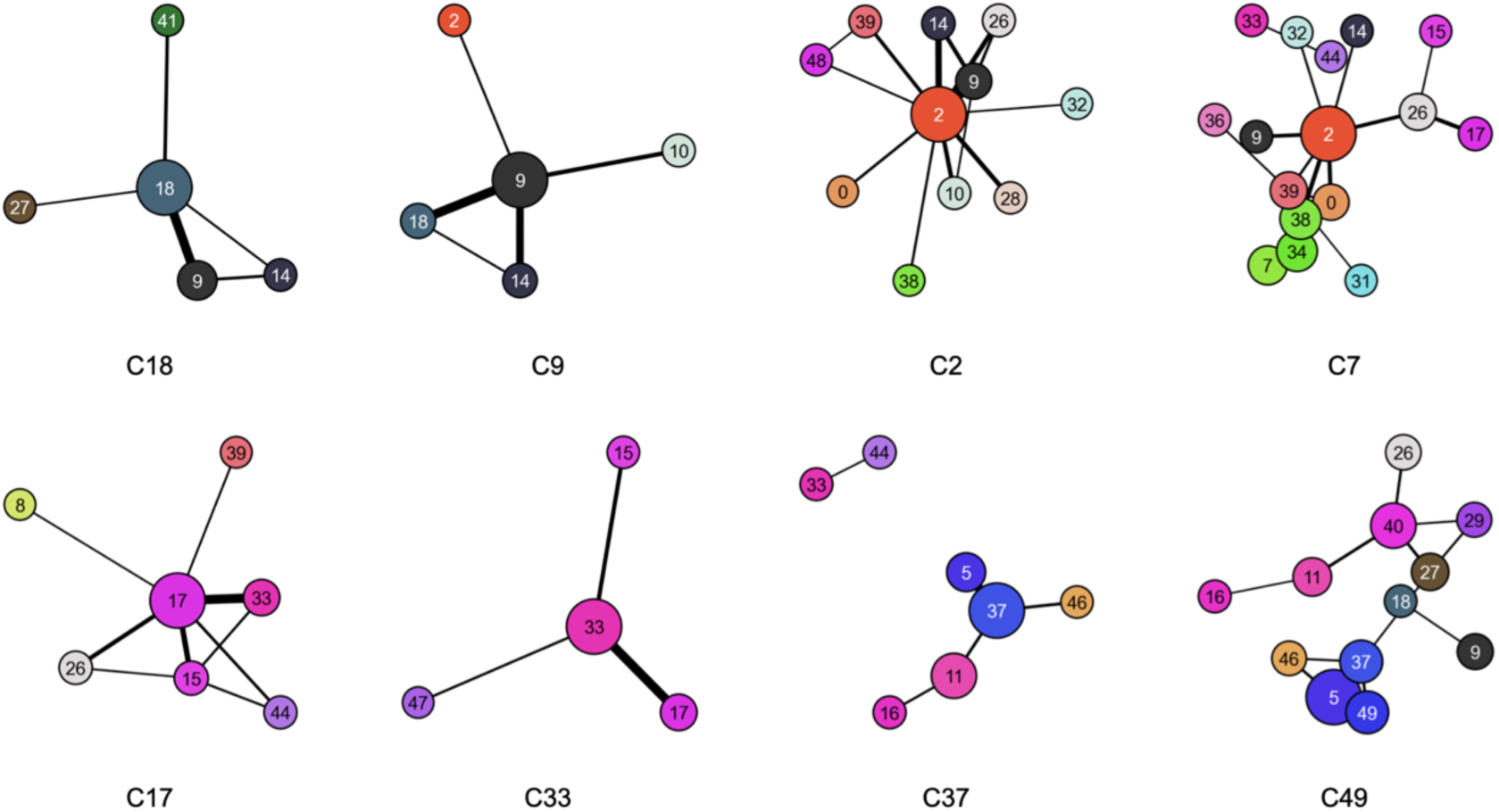
Signature adjacency in lung tumor biopsies. Adjacency graphs are constructed based on biopsies enriched for specific LTME signatures. Nodes and edges represent LTME signatures and their pair-wise interactions in selected biopsies.

**Supplementary Fig. 4:**
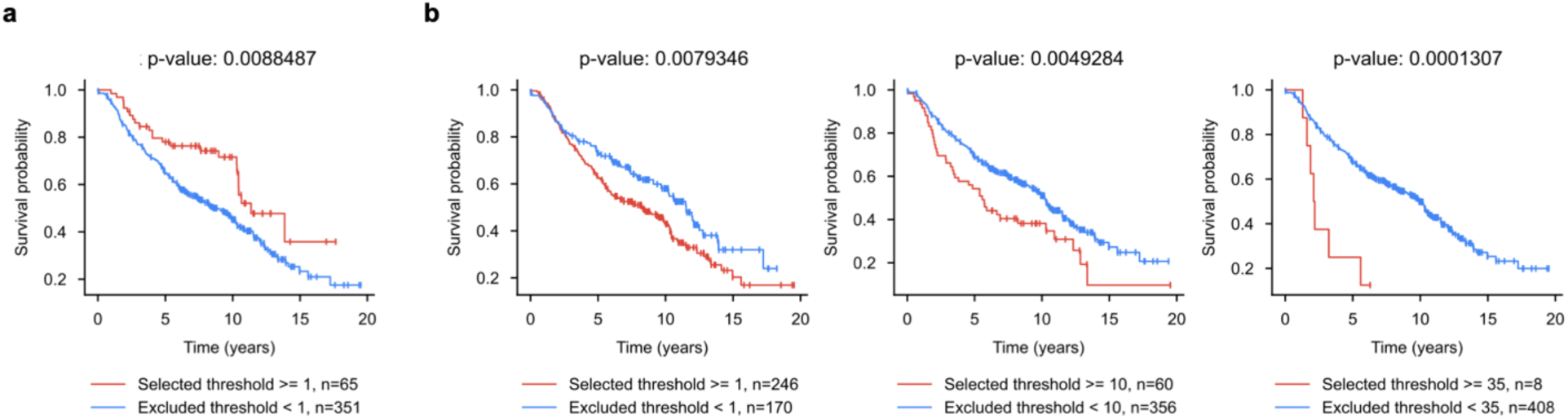
Survival analysis of patients with tumors enriched for specific signatures. a, Kaplan-Meier analysis on patients with C7-enriched tumor. **b**, Kaplan-Meier analysis on C10 enriched patients using LTME cutoff counts of 1, 10, and 35.

**Supplementary Fig. 5:**
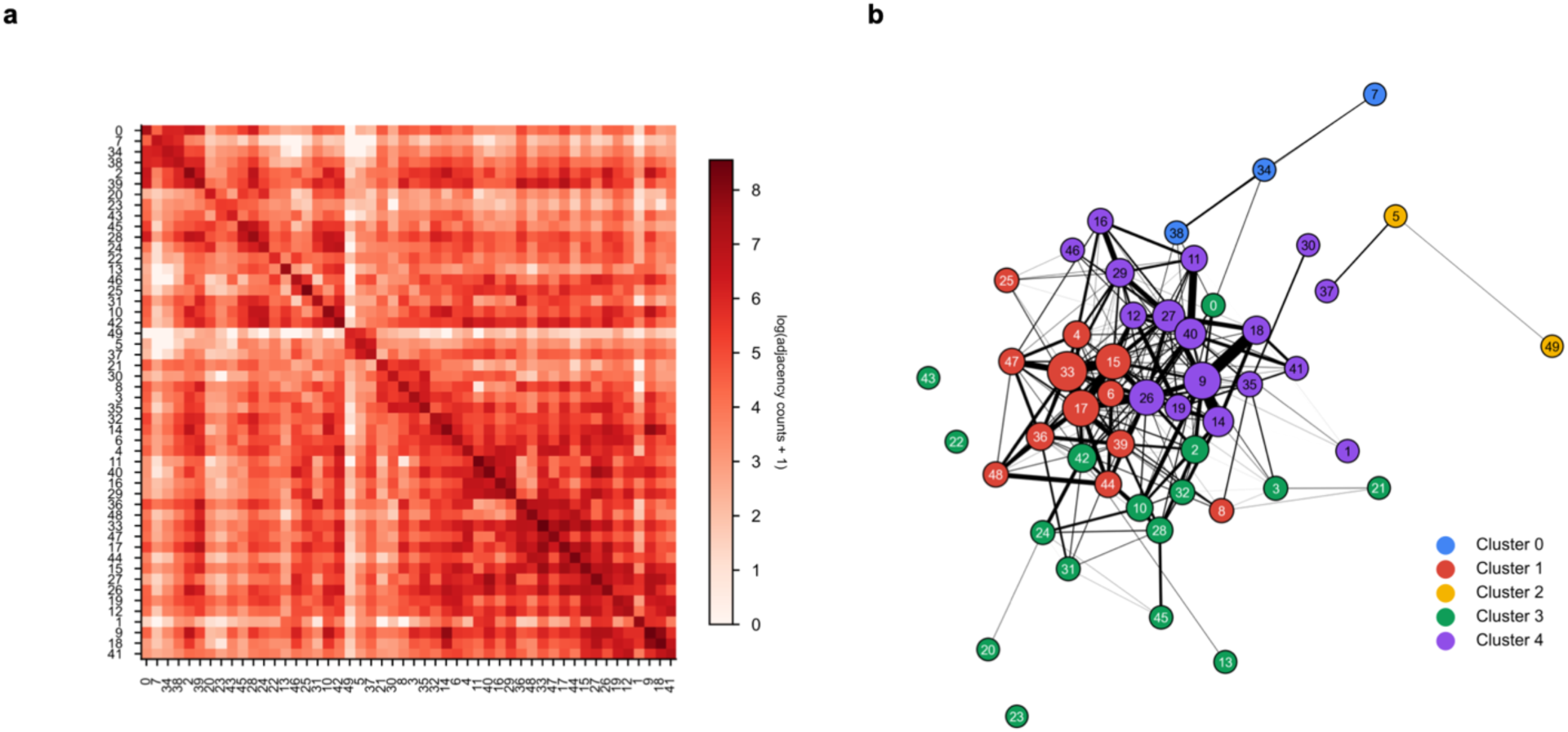
**Signature adjacency and spectral clustering. a**, Heatmap of signature adjacency matrix. An entry (i, j) in the matrix denotes the number of edges between two LTMEs from signature i and j. **b**, 5 clusters are generated from spectral clustering and used to color all 50 signatures and is used to color global adjacency graph.

**Supplementary Fig. 6:**
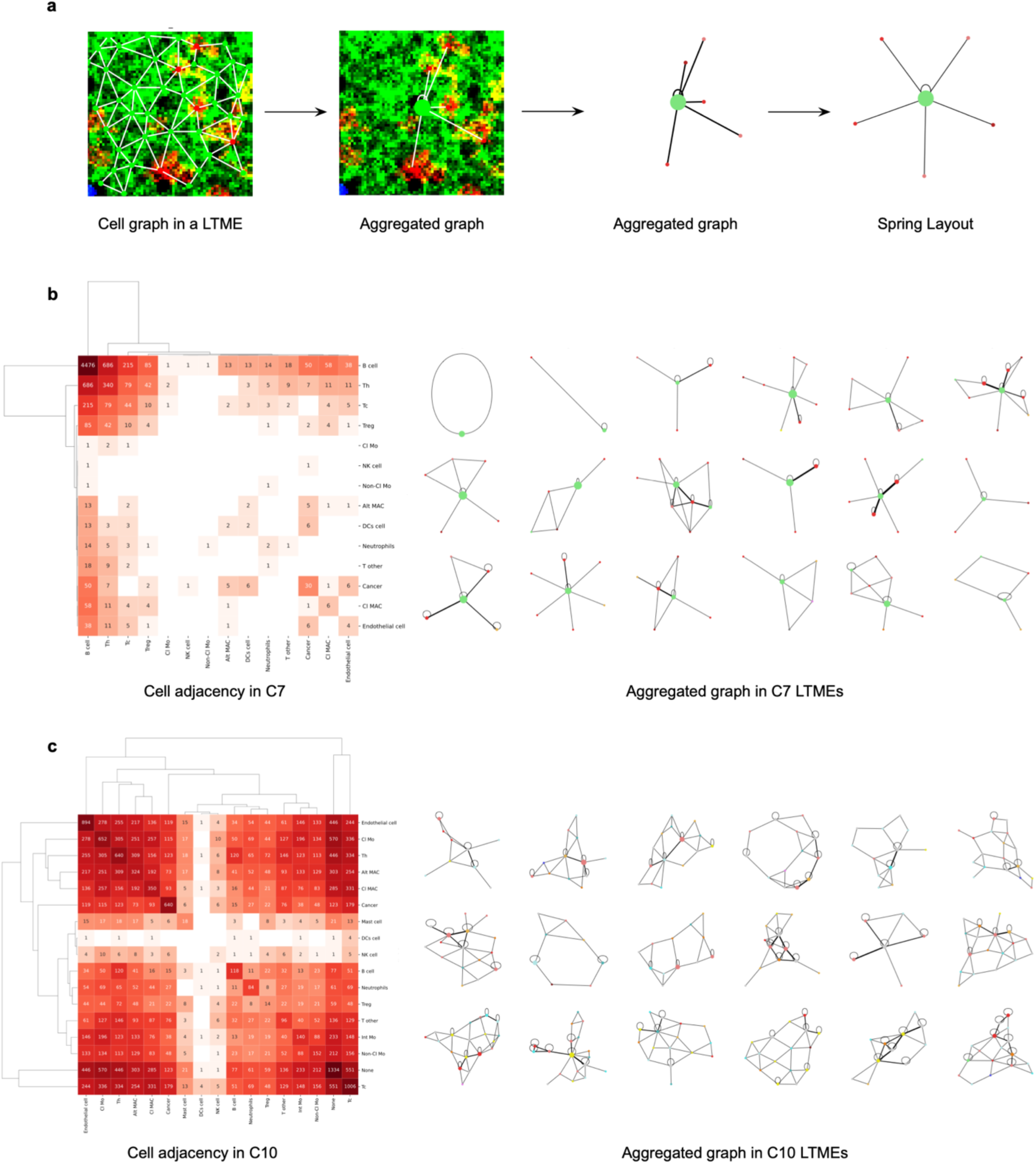
**Cell-to-cell interaction comparison between signatures C7 and C10. a**, A schematic showing conversion of an LTME with cell segmentations to a cell interaction graph. **b-c**, cell adjacency matrices aggregated across all LTMEs in C7 (**b**) and C10 (**c**) (left) and examples of LTME cell interactions after aggregation from C7 (**b**) and C10 (**c**) (right).

